# Bimodal Remapping of Visual Grids

**DOI:** 10.1101/2021.10.30.466568

**Authors:** Xiaoyang Long, Jing Cai, Bin Deng, Zhe Sage Chen, Sheng-Jia Zhang

**Affiliations:** Department of Neurosurgery, Xinqiao Hospital, Army Medical University, Chongqing 400037, China; Department of Psychiatry, Department of Neuroscience & Physiology, Neuroscience Institute, New York University School of Medicine, New York, NY 10016, USA

**Keywords:** grid cells, secondary visual cortex, remapping, bimodal grid patterns

## Abstract

Spatially modulated neurons from the rat secondary visual cortex (V2) show grid-like firing patterns during freely foraging in open-field enclosures. However, the remapping of the V2 grid cells is not well understood. Here we report two classes of V2 grid cell populations with distinct remapping properties: one regular class with invariant grid field patterns, and the other bimodal class that has remapping induced by environmental manipulations such as changes in enclosure shape, size, orientation and lighting in a familiar environment. The bimodal V2 grid cell pattern remains stable regardless of the follow-up manipulations, but restores to the original firing pattern upon animal’s re-entry into the familiar environment on the next day or from the novel environment. The bimodal V2 grid cells are modulated with theta frequency during the course of remapping and stabilize quickly. We also found conjunctive bistable V2 grid cells with invariant head directional tuning. Overall, our results suggest a new grid cell mechanism in V2 that is different from the medial entorhinal cortex (MEC) grid cells.

**Highlights:**

- Bistable V2 grid cells display bimodal or bistable remapping
- V2 grid cell firing patterns are not disrupted in darkness
- V2 grid cells preserve theta frequency modulation during remapping

## Introduction

Remapping of neuronal receptive fields reflects neural plasticity in response to changes in local or global environment, task context or stimulus cue (Latuske et al., 2017). Grid cells in the medial entorhinal cortex (MEC) is known to display spatial firing fields with a grid-like pattern (Hafting et al., 2005). Grid cells may change their firing patterns following environmental changes, which often involve translation and rotation of fields, but not much in rate-remapping fashion as reported in hippocampal place cells (Hafting et al., 2005, Fyhn et al., 2007, Barry et al., 2007, Jeffery, 2011, Barry et al., 2012, Marozzi et al., 2015). In humans, grid-like responses have been reported outside of the hippocampal formation in fMRI experiments [for a review, see (Raithel and Gottfried, 2021)], including the orbitofrontal cortex (OFC) (Constantinescu et al., 2016), prefrontal cortex (PFC) (Bao et al., 2019, Doeller et al., 2010) and anterior cingulate cortex (ACC) (Jacobs et al., 2013). Until very recently, grid cells have been discovered in the primary somatosensory cortex (S1) (Long and Zhang, 2021) and the secondary visual cortex (V2) from freely foraging rats in two-dimensional environmental enclosures (Long et al., 2021). However, the remapping of those sensory grid cells have not been systematically investigated, the potential circuit mechanisms of underlying the partial or global remapping of grid cells remains elusive.

Here we report two distinct subclasses of recorded V2 grid cells, which displayed two different stability in remapping patterns during environment changes in enclosure, one showing invariant grid field patterns and the other displaying translation and rotation in grid field patterns. Importantly, this bimodal remapping of V2 grid cells was not shown in the grid cells of rat MEC or S1, and was not found in other simultaneously recorded V2 spatially tuned cells (such as V2 place cells, V2 head-direction cells and V2 border cells). Additionally, unlike MEC grid cells that have disrupted firing patterns in the absence of visual input (Chen et al., 2016), the bimodal V2 grid cells have stable firing field independent of the visual input. MEC grid cells are reported to have modular grid scales and theta (6-12 Hz) modulation (Barry et al., 2012, Stensola et al., 2012, Deshmukh et al., 2010, Hafting et al., 2008). We found similar theta modulation among V2 grid cells, but did not observe any modular grid structure that might be due to sparse sampling. Moreover, theta frequency did not change during darkness where the visual input was absent. Thus, we first identify V2 bimodal/bistable grid patterns, which are distinct from firing properties of MEC grid cells.

## Results

### Bimodal remapping of V2 grid cells

From a total of 70 V2 grid cells of freely foraging rats identified in our previous study (Long et al., 2021), we began by asking whether V2 grid cells retained their spatial firing patterns in a similar manner as MEC grid cells did when animals responded to manipulations of distinct environments (Hafting et al., 2005, Fyhn et al., 2007, Barry et al., 2007, Barry et al., 2012). In our experimental design, we recorded and calculated only 40 out of 70 identified V2 gird cells (*n* = 6 rats) systematically for at least 20 min from freely foraging rats in open two-dimensional enclosures. The recordings were continued for four consecutive days consisting of several different sessions daily, each with unique experimental manipulation and separated by an overnight sleep in a home cage (**Fig. 1a**). Surprisingly, contrary to the context-invariant property of classical MEC grid cells, we observed that a proportion of V2 grid cells (15/40 = 37.5%) switched to another distinct spatial firing pattern upon entry into manipulations of the changed environment, continued to retain its switched spatial firing patterns and did not revert back to its original spatial firing pattern across a series of environmental feature manipulations (**Fig. 1**). We therefore named this class of V2 grid cell as the bimodal V2 grid cell.

**Figure 1.**
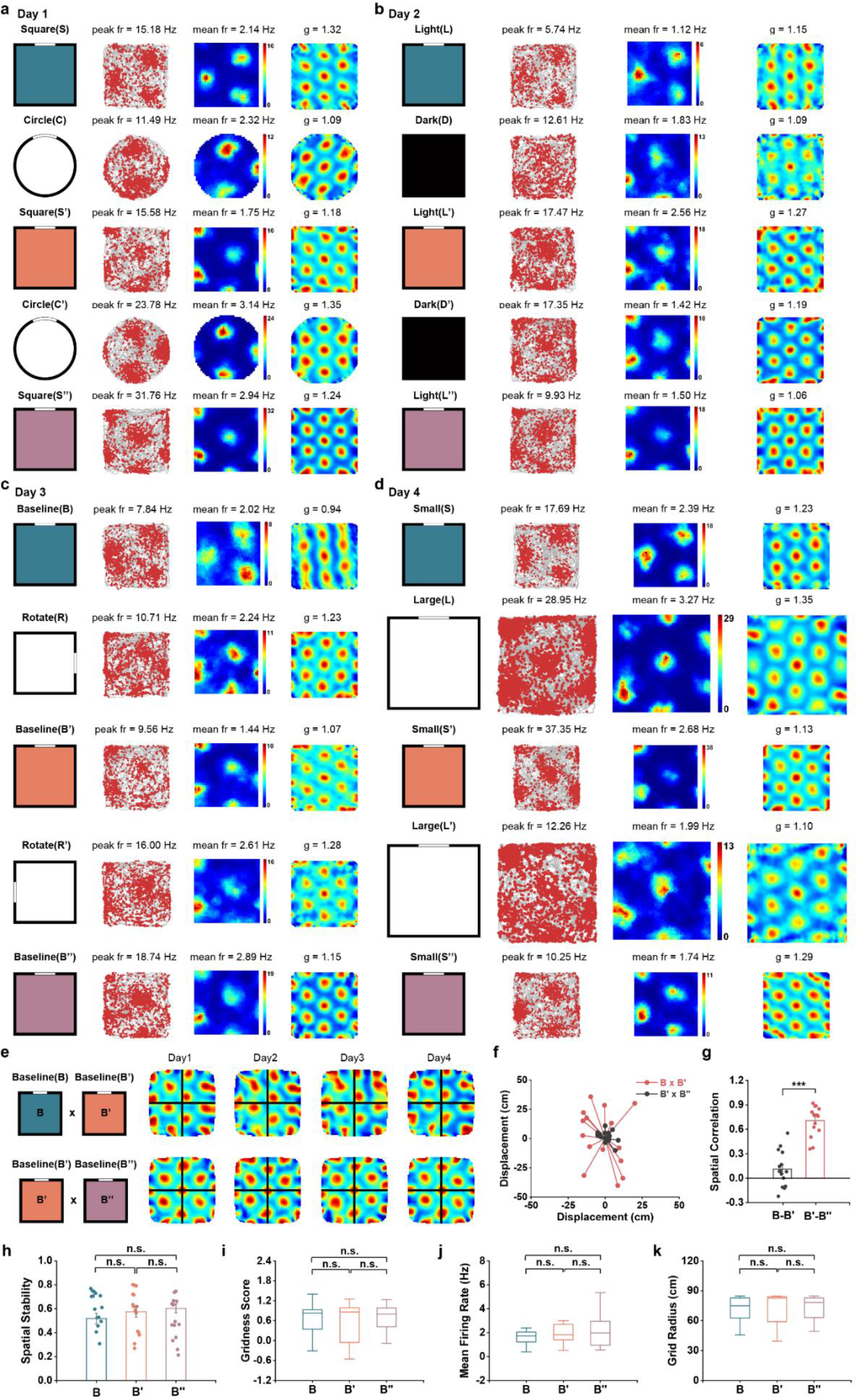
Bimodal remapping of firing patterns in V2 grid cells triggered by experimental manipulations. (a) Remapping of V2 grid firing patterns in different shapes of running boxes. (b) Remapping of V2 grid firing patterns after total darkness. (c) Cue triggered bimodal remapping of V2 grid firing patterns. (d) Remapping of grid firing patterns in different sizes of running boxes. (e) Cross-correlogram for the same cell during the same baseline condition after remapping (B versus B’) or after stabilization (B’ versus B’’). The peaks of the cross-correlogram were offset from the origin after environment change. (f) Vector diagram showing the grid displacement from the center of the cross-correlogram to the nearest peak. (g) Spatial correlation between two baselines after environmental manipulation (B-B’) and after the switch of grid firing patterns (B’-B’’). (h) Grid firing patterns were stable. (**i-k**) Gridness score, mean firing rate and grid scale showed no significant change during bimodal remapping.

For the first manipulation of environmental shape, we compared the spatial firing patterns of V2 bimodal grid cells across square (S)-circle (C)-square (S’)-circle (C’)-square (S’’) boxes in five 30-min recording sessions. The first, third and fifth session were within a familiar square enclosure and the second and forth were within a novel circular enclosure in the same recording room (**Fig. 1b**). Specifically, bimodal remapping was evoked by replacing the square box with the circle one (**Fig. 1b**). Like MEC grid cells, the spatial firing pattern of the representative V2 bimodal grid cell was realigned by the circular arena (second row in **Fig. 1b**). However, the V2 grid pattern did not revert to the original spatial structure (S) from the novel circle (C) back to the familiar square (S’) box, in contrast with the retained stable spatial alignment for MEC grid cells (Fyhn et al., 2007). When V2 grid cells were subsequently recorded in the circle (C’) and square (S’) arenas, the grid pattern still retained its bimodal firing pattern. After the completion of five consecutive recording sessions on day 1, recorded animals were returned to the home cage for sleep.

Notably, when animals returned to the familiar environment from home cage on day 2, the switched grid patterns of V2 bimodal grid cells reverted back to its original firing pattern observed during the first recording on day 1 (the first row in **Fig. 1c**). However, the switched grid patterns emerged again in the total darkness (D) and remained constantly stable during consecutive light-darkness-light (L-D-L’) sessions in the same familiar recording room. These results indicated that this bimodal grid pattern was not disrupted in the absence of visual inputs.

Again, the shifted grid spatial firing patterns changed back to its original firing structure on the first recording session on day 3. We further examined whether rotating the cue card can evoked bimodal remapping of V2 grid cells (**Fig. 1d**). The clockwise 90° rotation (R) of the cue card evoked bimodal switch from the baseline (B), and the remapped grid structure did not reverse to its original firing pattern once back to the baseline condition (B’), in contrast to the MEC grid cell remapping during the similar cue-rotation manipulation (Fyhn et al., 2007). Similarly, after counterclockwise 90° rotation (R’) of the cue card, the bimodal grid vertices remained similar between rotation (R’) and baseline (B’).

On day 4, we examined whether bimodal remapping could be induced in a larger environment from the baseline trial (the first row in **Fig. 1e**). Again, we found similar remapping patterns in the recorded V2 grid cells in a 1.5 x 1.5 m^2^ novel square box (L) and remained stable when back to the standard 1 x 1 m^2^ recording enclosure (S). The bimodal grid structure stayed unaltered although the size of the running boxes was reduced back to 1 x 1 m^2^ (S’) again.

In contrast to the bimodal V2 grid cells, the majority of recorded V2 grid cells (25/40 = 62.5%) did not switch their spatial firing patterns after environmental change in the same familiar room (**Fig. S1**), and we called this subpopulation as the regular V2 grid cells. To compare the bimodal with regular V2 grid cells, we calculated the mean displacement of the peak of the spatial cross-correlogram between the original baseline (B) and the baseline (B’) after environmental change. We found that the mean displacement of regular V2 grid cells was centered on the origin with the mean displacement being 5.25 ± 0.11 cm (mean ± s.e.m.), which was significantly smaller than that for bimodal V2 grid cells (**Fig. S1f**, *Z* = 4.88, *P* = 1×10^-6^, two-tailed Mann-Whitney U test, *n* = 15 and 25). In turn, the spatial correlation coefficient was significantly higher in V2 regular grid cells than in V2 bimodal grid cells (**Fig. S1g**, *Z* = 4.57, *P =* 5.0×10^-6^).

As a comparison, we independently recorded 15 MEC grid cells from two new implanted rats under the same familiar environment (**Fig. S2**). However, we found that the MEC grid cell did not exhibit bimodal remapping patterns. Both spatial correlation coefficient (**Fig. S2g**, *Z* = 4.13, *P* = 3.7×10^-5^, two-tailed Mann-Whitney U test, *n* = 15 and 15) and displacement of peaks of cross-correlogram (**Fig. S1f**, *Z* = 4.34, *P* = 1.5×10^-5^) were significantly different from those of V2 bimodal grid cells.

To quantify the displacement of bimodal grid patterns in V2, we computed the cross-correlograms between three baseline sessions. The peak of the cross-correlograms (B×B’) was offset from the origin (first row of **Fig. 1e**), and the mean displacement was 22.72 ± 2.54 cm (red dots in **Fig. 1g**). However, when comparing B’ and B’’ sessions, the peak of the cross-correlograms was significantly smaller (two-sided Wilcoxon signed-rank tes, *Z* = 3.41, \*\*\**P* = 0.001, *n* = 15), centered on the origin (the second row in **Fig. 1e**) with the mean displacement being 6.05 ± 0.78 cm (black dots in **Fig. 1f**). This observation indicated that V2 grid cells re-aligned after bimodal remapping of spatial firing patterns. Moreover, the spatial correlation in B’×B’’ was significantly higher than that in B×B’ (**Fig. 1g**, *Z* = 3.41, \*\*\**P* = 0.001, *n* = 15). Additionally, spatial stability was preserved among three baseline conditions (**Fig. 1h,** t, B-B’: *Z* = 1.02, *P* = 0.31; B’-B’’: *Z* = 0.8, *P* = 0.43; B-B’’, *Z* = 1.65, *P* = 0.1). We further compared the grid firing properties of V2 bimodal grid cells, gridness score, mean firing rate and grid scale among three baseline sessions (**Figs. 1i-k**) and found no significant difference between them (two-sided Wilcoxon signed-rank test, gridness score: B-B’: *Z* = 0.17, *P* = 0.86; B’-B’’: *Z* = 1.53, *P* = 0.13; B-B’’, *Z* = 0.34, *P* = 0.73; mean firing rate: B-B’: *Z* = 0.68, *P* = 0.5; B’-B’’: *Z* = 0.4 5, *P* = 0.65; B-B’’, *Z* = 0.68, *P* = 0.5; grid radius: B-B’: *Z* = 0.06, *P* = 0.96; B’-B’’: *Z* = 0.85, *P* = 0.39; B-B’’, *Z* = 0.17, *P* = 0.87). Together, these results suggested that V2 grid cells exhibited intrinsic bimodal firing patterns; the environmental manipulation in the familiar recording room could evoked bimodal remapping in a subpopulation of V2 grid cells; and the V2 grid-like firing structure became stabilized after remapping.

### Restoration of V2 bimodal grid patterns upon re-entry into the familiar environment

We further investigated whether bimodal remapping of V2 grid cells are reversible in their firing patterns, and recorded another bimodal V2 grid cell (with conjunctive head-direction tuning) in both familiar and novel environments across multiple days (**Fig. 2**). We first triggered the bimodal remapping of the V2 grid cell from the original firing pattern in the standard square (S) box to a new firing pattern in the circle (C) box in the same familiar room (**Fig. 2a**). To reset grid firing pattern, we then conducted a series of recordings starting from the familiar room (S’) to another novel room with distinct distal landmarks (the fourth row in **Fig. 2a**). Consistent with the substantial realignment of grid cells in MEC, V2 bimodal grid cells showed a significant shift in grid structure in the novel environment. However, the grid pattern reversed back to its original firing structure upon animal’s re-entry into the square box (S’’) in the familiar room. Interestingly, the bimodal firing pattern can be repeatedly induced upon changes in two different environmental geometries (C’ and S’’’ enclosures in **Fig. 2a**). Notably, the head directional tuning of the conjunctive V2 bimodal grid cell switched both directional tuning and bimodal grid pattern in the novel room (the fourth rows in **Fig. 2a** and **Fig. 2b**), indicating that the conjunctive V2 bimodal grid cells maintained a coherent ensemble for both head directionality and grid structure in response to manipulations of environmental feature.

**Fig. 2.**
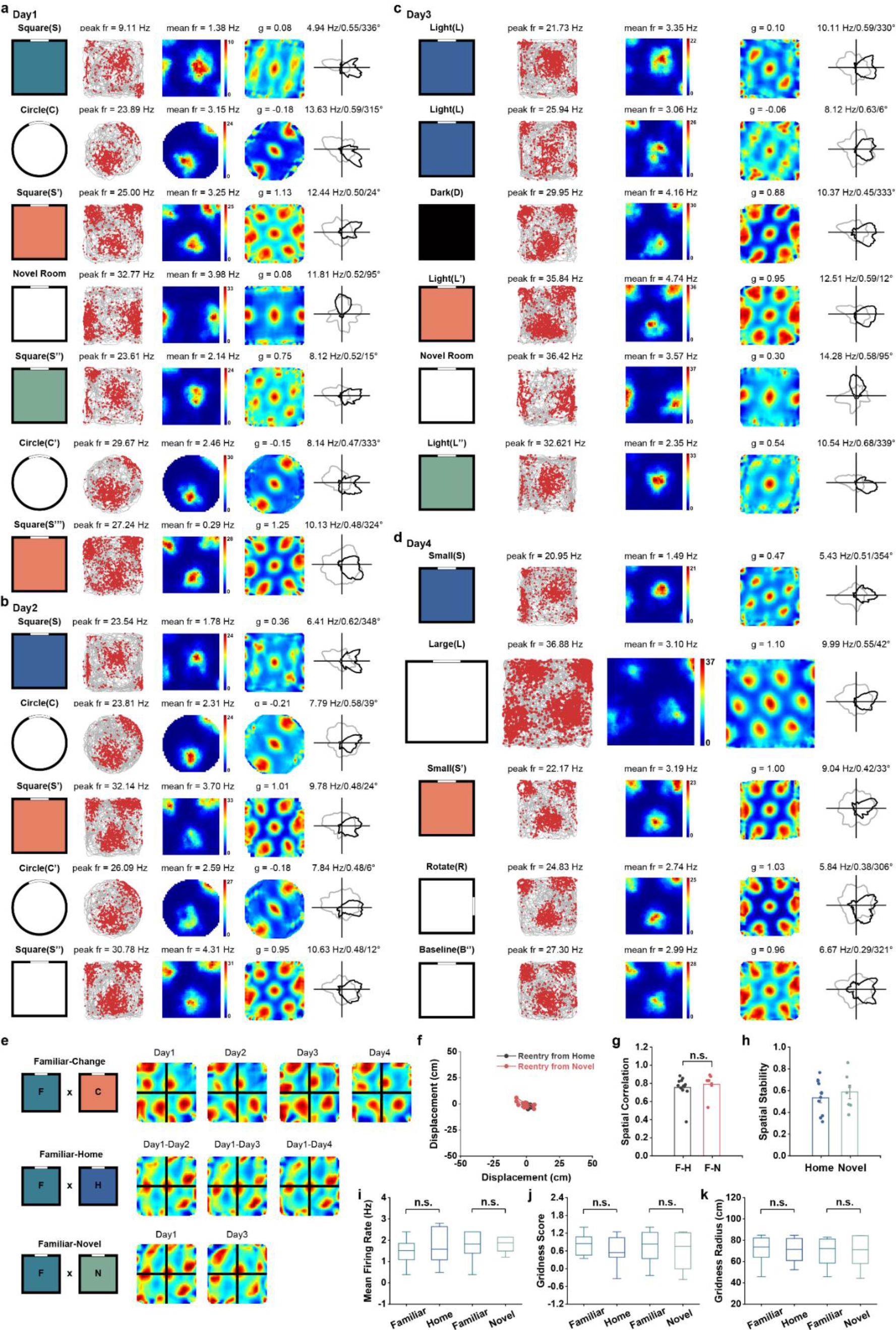
V2 grid cell grid firing pattern stabilized after bimodal remapping and restored upon animal’s re-entry into the same familiar environment. (a) Spatial responses of a representative V2 conjunctive grid cell showing bimodal remapping of firing pattern after environmental change and restoration of grid firing pattern upon animal’s re-entry from another room into the same familiar environment. (b) The same grid cell showing stabilization of grid firing pattern after bimodal remapping and did not change firing pattern in a different shape of running boxes, which led to bimodal remapping of firing patterns in the first place. (c) The firing patterns of grid cell remained stable when the environment was not changed and reset of grid pattern after animal’s re-entry from another novel environment. (d) The firing pattern of the same grid cell was reset after being brought from home cage into the familiar room. (e) Cross-correlogram for the same cell between the baseline condition in the familiar environment and the same baseline condition after environmental change (upper row), the same baseline condition after re-entry into the familiar room from home cage (middle row) and the same baseline condition after animal’s re-entry into the familiar room from a novel environment (bottom row). The peaks of the cross- correlogram were centered at the origin after re-entry. (f) Vector diagram showing the grid displacement from the center of the cross-correlogram to the nearest peak. (g) Distribution of spatial correlation coefficients between two baselines after re-entry either from the home cage or a novel room. (h) Spatial stability between the baseline condition in the familiar environment and the same baseline condition after re-entry into the familiar room from home cage and the same baseline condition after re- entry into the familiar room from a novel environment. (**i**-**k**) Gridness score, mean firing rate and grid scale were stable before and after animal’s re-entry into the same familiar environment.

Could bimodal grid patterns reset after returning to the home cage, equivalent to a possible reset of the path integrator? To address this possibility, we recorded the same V2 bimodal grid cell from the home cage on day 2 and found that the original grid pattern emerged again (the first row in **Fig. 2b**). Subsequent box shape-change caused the appearance of the bimodal grid patterns between the circle (C) and the familiar square running box (S’). The bimodal grid pattern was maintained despite further environmental manipulations (C’ and S’’ in **Fig. 2b**). In all box-shape manipulations within the same familiar recording room, the directionality of the recorded conjunctive V2 grid cell remained similar head directional tuning (the fifth columns in **Fig. 2b** and **Fig. 2d**).

We next asked if the original grid pattern of the bimodal grid cell was stable in the darkness without any change in the box-shape. As expected, the grid-like firing pattern was stable between two repeated light-on baseline sessions (the first and the second rows in **Fig. 2c**). However, the light-off caused the sudden shift in the grid structure (the third row in **Fig. 2c**), and darkness-induced bimodal grid structure remained stable in the following light condition (L’) in the familiar square enclosure. Notably, the bimodal grid pattern was switched again to a distinct grid structure in another novel recording room (the fifth row in **Fig. 2c**), with its conjunctive head directional tuning rotated (the fifth columns in **Fig. 2c**). Interestingly, the novel environment-induced grid firing pattern was completely reversed back to its original firing mode after re-entry into the familiar room (the sixth row in **Fig. 2c**).

On day 4, we further verified the bimodal remapping in a larger environment (**Fig. 2d**). As predicted, the bimodal grid pattern was evoked upon animal’s entry into a larger enclosure (L) from the small box (S). After returning to the small recording box (S’), the grid pattern was transformed into another firing mode (the third row in **Fig. 2d**). Further manipulation of the local cue in the standard running box (R) imposed no difference in terms of its grid mode (the fourth row in **Fig. 2d**). The grid mode was preserved despite the cue card being rotated to its original orientation (the fifth row in **Fig. 2d**), suggesting that once the bimodal pattern was switched, further environmental change could not induce any substantial shift in its grid structure. We found that the peaks of the cross-correlograms for sessions before and after environmental change (F×C) were offset from the origin (the first row in **Fig. 2e**). However, on session before and after re-entry to the home cage, the peaks of the cross-correlograms (F×H) remained in the center (the second row in **Fig. 2e**) with the mean displacement being 4.92 ± 0.76 cm (black dots in **Fig. 2f**). Accordingly, the peaks of the cross-correlograms (F×H) were also centered on the origin before and after re-entry from the novel room (the third row in **Fig. 2e**) with the mean displacement being 3.07 ± 0.68 cm (red dots in **Fig. 2f**).

Furthermore, the spatial correlation between two baselines after re-entry either from the home cage or a novel room showed no significant difference (**Fig. 2g**, two-tailed Mann-Whitney U test, *Z* =1.27, *P* = 0.23, *n* = 12 and 7). Additionally, spatial stability after re-entry either from the home cage or a novel room was preserved (**Fig. 2h**, *Z* = 1.44, *P* = 0.17). To compare the grid feature of V2 bimodal grid cells during the reset, we calculated the gridness score, mean firing rate and grid scale among three baseline sessions and found no significant difference during reset (two-sided Wilcoxon signed-rank test, **Fig. 2i**, mean firing rate: Familiar-Home: *Z* = 1.18, *P* = 0.24; Familiar-Novel: *Z* = 0.34, *P* = 0.74; **Fig. 2j**, gridness score: Familiar-Cage: *Z* = 1.33, *P* = 0.18; Familiar-Novel: *Z* = 1.01, *P* = 0.31; **Fig. 2k**, grid radius: Familiar-Home: *Z* = 0, *P* = 1.00; Familiar-Novel: *Z* = 0, *P* = 1.00). Importantly, the bimodal remapping of grid firing patterns was not induced or accompanied by shift of cluster diagram or spike waveform (**Fig. S10**). Of note, we found no significant change in either the separation of clusters (**Figs. S10c-d**), spike amplitudes (**Figs. S10e-f**) or spike widths (**Figs. S10g-h**) in various comparisons. Together, the bimodal V2 grid firing patterns can be restored upon reentry into the same environment.

### Characterization of rotation and deformation in V2 bimodal grid patterns

What is the difference between two distinct modes of V2 grid firing pattern? Consistent with MEC grid cells (Krupic et al., 2015, Stensola et al., 2015), the inner six fields of the auto-correlogram of V2 grid cells formed an ellipse (**Fig. 3a**) instead of a perfect circle. To quantify the change in spatial firing pattern for V2 bimodal grid cells, we fitted an ellipse to the inner hexagon of auto-correlogram (**Fig. 3b**) and defined semi-major and semi-minor axes (**Fig. 3c)**. The orientation of the fitted ellipse was defined as the angle between the horizontal axis and semi-major axis. We compared the elliptification and orientation of spatial patterns under three conditions: in the familiar room (F), after the environmental change (C), and re-entry into the same familiar room (R). As shown in two V2 grid cell examples, the grid pattern was deformed and rotated after the environmental change (**Figs. 3a-b**), with almost complete overlapping between familiar and re-entry trials. We further found significant difference in elliptic orientation (**Fig. 3e**, two-sided Wilcoxon signed-rank test, *Z* = 3.07, \*\**P* = 0.002, *n* = 15), reflecting a clear deformation of grid patterns after environmental change. Additionally, grid orientation also varied significantly after bimodal remapping (**Fig. 3f**, *Z* = 2.73, \*\**P* = 0.006). However, the degree of elliptification (“ellipiticity”) was preserved (**Figs. 3g-h**, Familiar-Change: *Z* = 0.63, *P* = 0.53; Change- Re-entry: *Z* = 0.97, *P* = 0.33; Familiar-Re-entry: *Z* = 0.34, *P* = 0.73), suggesting that the distortion of the hexagonal symmetry of grid patterns maintained and underwent non-coaxial rotation between two distinct V2 grid cell firing modes.

**Fig. 3.**
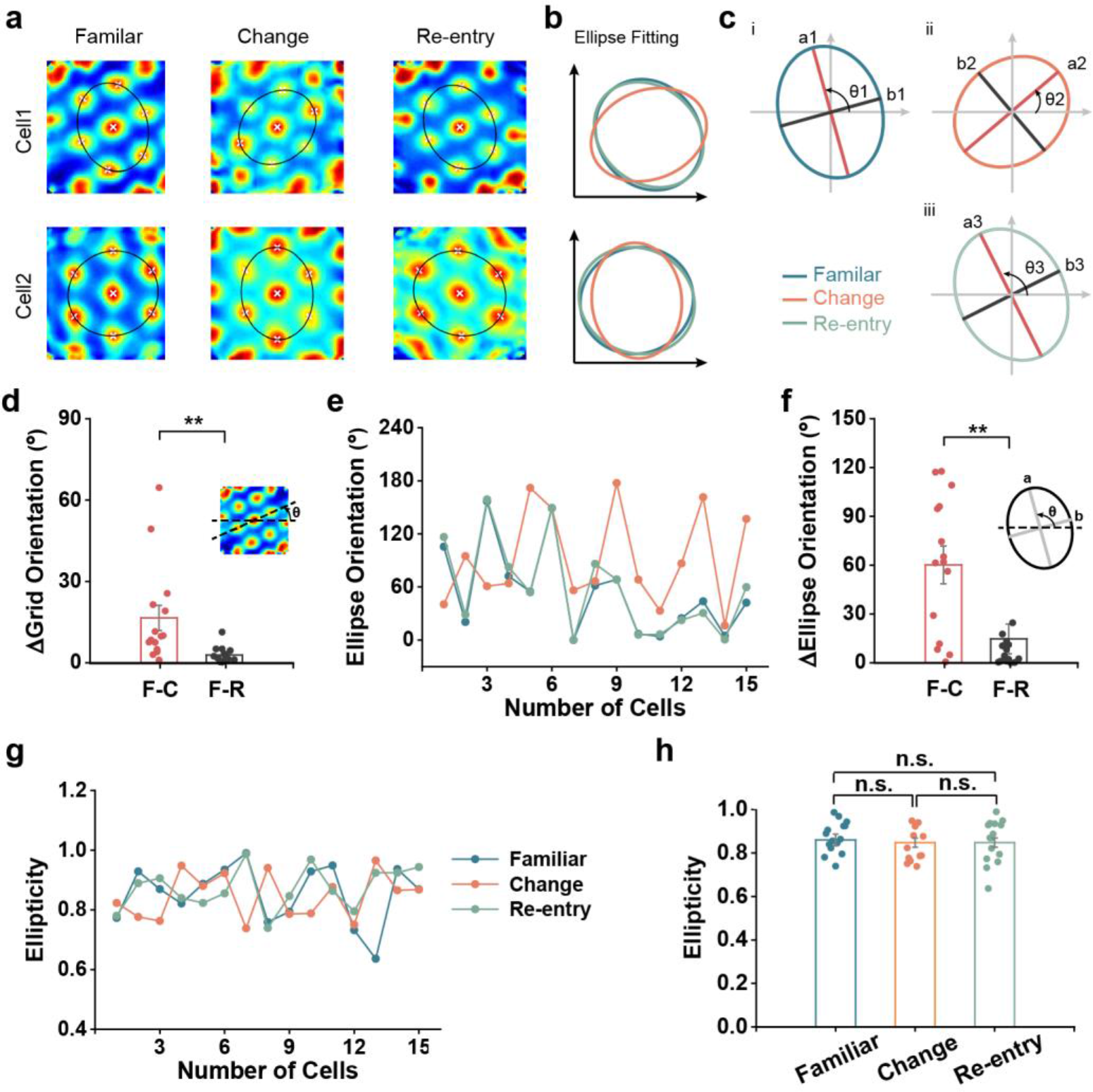
Grid realignment by bimodal switch reflects shearing. (a) Representative ellipse fitting of auto-correlogram of two visual grid cells in the familiar environment (left panels), back to the same baseline condition after environmental change (middle panels) and re-entry into the same familiar environment (right panels). (b) Deformation of grid firing patterns during transformation between the bimodal firing patterns. The ellipse deformation restored after re-entry to the same familiar environment. (c) Diagram illustrating the calculation of ellipse orientation and the axes used to calculate ellipticity in the three conditions above. (d) Differences in grid orientation between a familiar environment and environmental change (F-C) were significantly higher than that between a familiar environment and re-entry to the same environment (F-R). (e) Distribution of the horizontal orientations of the fitted ellipses for the three conditions above. (f) Orientation difference between a familiar environment and environmental change (F-C) was significantly higher than that between first entry and re-entry to the same environment (F-R). (g) Distribution of the ellipticity of the fitted ellipses for the three conditions. (h) Ellipticity of the grid patterns showed no significant difference by bimodal remapping.

Since elliptification of the grid cells reflects shearing along the environmental boundaries, we next sought to identify the angular offset of the grid axes to the box axes. In a perfect hexagonal grid cell, a rotation of 7.5°, 22.5°, or 37.5° would result in asymmetrical distribution from the environmental boundary (**Fig. 4a**). To quantify the degree of the rotational offset, we identified three grid axes with the minimal angular deviation from the nearest wall (**Fig. 4b**). The minimal value of the angular offset of grid axes from the walls for MEC grid cells was near 7.5°, which reflected the possible shear force arising from anchoring to the geometrical borders and thus minimized symmetry with the environmental boundaries (Stensola et al., 2015). Therefore, we further investigated if the asymmetrical relationship to the environment boundaries had been reshaped during the transition to bimodal remapping. We found that despite obvious deformation of grid patterns during bimodal remapping, the minimal angular deviation of the grid axes from the walls was maintained (**Fig. 4c**). The minimal angular offset exhibited a similar distribution after bimodal remapping (**Fig. 4d**), and the mean absolute angular offset was not significantly different (**Fig. 4e**, two-sided Wilcoxon signed-rank test, Familiar-Change: *Z* = 0.97, *P* = 0.33; Change-Reentry: *Z* = 0.54, *P* = 0.59; Familiar-Reentry: *Z* = 1.16, *P* = 0.25, *n* = 15). The mean absolute angular deviation was 9.95 ± 1.74° in familiar environment, 7.26 ± 1.58° after remapping and 8.96 ± 1.92° back to the same familiar room, not far from the 7.5° angular offset that would have minimized symmetry or overlap with the cardinal and diagonal axes of the environment. In summary, the rotational offset of grid patterns remained consistent despite dramatic rotation and distortion during bimodal V2 grid cell remapping, suggesting that the asymmetric relationship with the environmental boundary was preserved in the two distinct V2 grid cell firing modes.

**Fig. 4.**
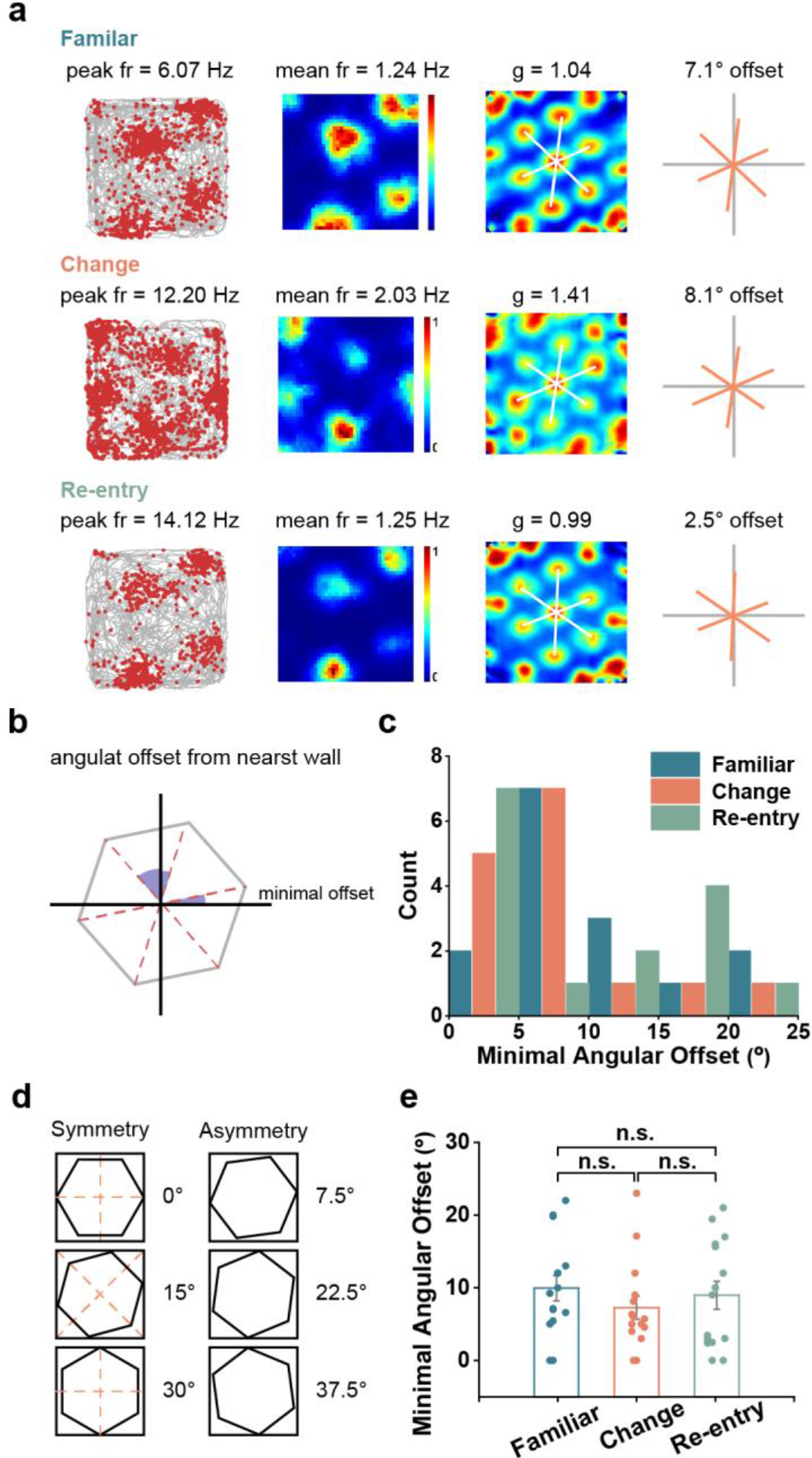
Grid alignment after stabilization remained asymmetric with respect to the cardinal and diagonal axes of the environment. (a) Representative grid cells in a familiar environment (upper panels), after environmental change (middle panels) and re-entry (bottom panels). For each condition, three grid axes were plotted in the right. (b) Schematic illustrating the definition of angular offset from the boundaries of the environment. (c) Distribution of the smallest angular offset of grid axes from the nearest wall. Under the three conditions, the orientations of grid axis with minimal offset from the box walls were centered around 7.5° (red vertical line). (d) Diagram illustrating the rotation of grid firing pattern in different angles resulting the symmetric and asymmetric alignment of the external environment. Red dashed line showing the axes of symmetry. (e) Minimal angular deviation of grid axes from the nearest wall did not differ significantly among three conditions.

### V2 bimodal grid patterns only switch in the familiar environment

We further determined whether bimodal remapping of V2 grid cells also occurred in a novel environment after recording in the familiar room first. To do so, we recorded the spatial responses of the representative bimodal grid cell from **Fig.1** in two novel rooms, each with distinct distal landmarks (**Fig. S3**). The bimodal grid cell was first recorded between small-large-small (S-L-S’) running box sessions on day 1 (**Fig. S3a**), and the firing patterns between two small boxes remained the same. On day 2, the grid cell was recorded across square-circle-square (S-C-S’) sessions (**Fig. S3b**), with the grid structure maintained between two square boxes. To further validate the unchanged property of V2 bimodal grid cells in the novel environment, on day 3 we recorded the same grid cell in another novel room and found that the V2 grid cell firing pattern did not shift in response to environmental changes across square-circle-square (S-C-S’) trials (**Fig. S3c**). This observation was confirmed by cross-correlogram between two baseline sessions (B×B’) with peaks almost centered at the origin (**Fig. S3d**). The finding suggested that V2 bimodal grid cells seemed to measure geometric changes only in the familiar environment.

### Preservation of theta modulation during remapping of V2 bimodal grid patterns

Theta oscillation (4-8 Hz) of hippocampal or cortical local field potentials (LFPs) is vital for formation of grid cells (Buzsáki and Moser, 2013). To explore the influence of the temporal oscillation on the spatial firing patterns of V2 bimodal grid cells, we measured the LFP and spikes simultaneously from the freely-foraging animals. Running was remarkably associated with strong theta rhythmicity in V2, and the majority of V2 bimodal grid cells (12/15 = 80%) displayed clear theta-modulation in their spike auto-correlogram (**Figs. S4a**-**d**). We observed continuous theta rhythmicity during the transition of grid bimodal firing patterns, and found no significant difference in both theta modulation index [**Fig. S4e,** two-sided Wilcoxon signed-rank test, *Z* = 0, *P* = 1.00 (B-B’), *Z* = 1.82, *P* = 0.07 (B-B’’) and *Z* = 1.39, *P* = 0.16 (B’-B’’), *n* = 15], and peak theta frequency [**Fig. S4f,** *Z* = 0.34, *P* = 0.73 (B-B’), *Z* = 0.79, *P* = 0.43 (B-B’’) and *Z* = 1.73, *P* = 0.08 (B’-B’’)]. These results suggested that the majority of V2 bimodal grid cells preserved temporal periodicity and theta modulation during the transformation of spatial firing patterns. Strong theta modulation was also observed during the re-entry into the same familiar environment which caused restoration of the original grid structures (**Figs. S5a-d**). We found no significant change in both theta modulation index (Familiar-Home: *Z* = 0.63, *P* = 0.53; Familiar-Novel: *Z* = 1.018, *P* = 0.31, two-sided Wilcoxon signed-rank test, *n* = 5 and 3) and peak theta frequency (Familiar-Home: *Z* = 1.75, *P* = 0.08; Familiar-Novel: *Z* = 0.94, *P* = 0.35).

### V2 bimodal grid patterns stabilize rapidly upon changes in the familiar environment

Does the bimodal grid structure stabilize rapidly or change gradually after the environmental change? To investigate this question, we split the 30-min recording session into three 10-min epochs, and found that the bimodal grid pattern was expressed instantly after the environment change (**Fig. S6a**). Firing fields appeared to be stable from the onset of the running session. The development of bimodal firing structures was estimated by correlating for each V2 bimodal grid cell and the whole session with five 2- min incremental blocks throughout the whole session. Notably, the spatial firing patterns were highly correlated during the course of recording time (**Fig. S6b**, Friedman’s test, *P* = 0.05), and the normalized mean firing rate was also stable during the course of recording (**Fig. S6c**, Friedman’s test, *P* = 0.27). Therefore, the bimodal remapping of V2 grid cells took place rapidly in response to manipulations in the familiar environment.

### Bimodal remapping of V2 grid firing patterns without disturbing head directionality

Due to the presence of diverse spatially-modulated V2 neurons (Long et al., 2021), one may wonder if the V2 bimodal grid cells arose from the local environment content or visual input, what had happened to other spatial cell types in V2 under the same experimental manipulations. Among 15 recorded V2 bimodal grid cells, we detected four V2 grid cells with conjunctive head direction tuning (**Fig. S7**). As expected, the grid firing pattern of the conjunctive bimodal grid cells preserved when there was no change in the familiar environment between the first and second baseline (L) sessions on day 1 (the first and second rows in **Fig. S7a**). Under the darkness (D) session, the spatial firing pattern was switched to another firing mode (the third row in **Fig. S7a**). The grid structure was stabilized when returning to light (L’) conditions (the fourth row in **Fig. S7a**) and did not change anymore in another follow-up baseline (L’) session (the fifth row in **Fig. S7a**). However, the head directionality of both conjunctive bimodal V2 grid cell and simultaneously recorded V2 head-direction cell was almost invariant throughout the whole recording session (the fifth columns in **Fig. S7a**).

To further validate the invariant head directionality of conjunctive bimodal grid cells, we recorded the same pair of V2 cells across square-circle-square (S-C-S’) sessions on day 2. Consistent with those switched grid patterns and preserved head directionality on day 1, we found different shapes of running boxes triggered the remapping of bimodal grid patterns (the first to the third rows in **Fig. S7a**) without perturbation of its head directionality (the fifth columns in **Fig. S7b**). In addition, the animal’s re-entry into the same familiar room from a novel environment (the fourth row in **Fig. S7a**) restored the grid firing pattern (the fifth row in **Fig. S7b**). To quantify the head directional tuning accompanying the bimodal remapping of V2 grid pattern, we measured the peak direction of head directional tuning of both conjunctive bimodal grid cells and co-recorded head-direction cells under three baseline conditions: in the familiar environment (B), after the environmental change (B’) and re-entry into the same environment (B’’), we detected no significant change in the peak direction of head direction tuning (**Fig. S7c**, Rayleigh test for uniformity: B-B’: median offset = 2.98°, *r* = 0.93, *P* = 1.43×10^-5^; B’-B’’: median offset = 2.98°, *r* = 0.96, *P* = 3.68×10^-6^; B-B’’: median offset = 5.95°, *r* = 0.90, *P* = 4.37×10^-5^).

Additionally, there was no significant difference in peak angular rate [**Fig. S7d**, two-sided Wilcoxon signed-rank test, B-B’: *Z* = 0.76, *P* = 0.45; B’-B’’: *Z* = 1.38, *P* = 0.17; B-B’’: *Z* = 0.56, *P* = 0.58] and mean vector length [**Fig. S7e**, B-B’: *Z* = 0.36, *P* = 0.72; B’-B’’: *Z* = 0.36, *P* = 0.72; B-B’’: *Z* = 0.71, *P* = 0.48]. It might be possible that V2 bimodal grid pattern and head directional tuning originate from different computational mechanisms. In other examples of simultaneously recorded units that included a V2 bimodal grid cell (**Fig. S9, i-iii**), we found a similar trend with bimodal mapping of grid patterns accompanying with invariant place or border spatial firing patterns during environmental changes. Finally, we compared the grid structure of simultaneously recorded bimodal grid cells during the environmental manipulations and determined whether ensembles of V2 bimodal grid cells remained coherent in remapping. Remarkably, two pairs of co-recorded bimodal V2 grid cells (one non-conjunctive grid cell and the other conjunctive grid cell) co-varied when the familiar environment changed, and then co-restored their original firing modes upon animal’s re-entry into the same familiar environment (**Fig. S6**).

## Discussion

The extrastriate (V2) cortex is the second major area in the visual cortex and the first region within the visual association area. V2 receives a primary feedforward input from V1 and lateral geniculate nuclei (LGN) and processes specific features of visual information (such as contour and shape); V2 projects topographically back to V1, and sends strong feedforward input to the upstreams of the visual cortex (V3, V4 and MT). In addition to the projections within the visual cortex, V2 also projects to three visual areas in the parietal cortex---the medial superior temporal (MST), parieto-occipital (PO) and ventral intraparietal (VIP) areas. Projections from the peripheral field representation of V2 of parietal areas could provide a direct route for rapid activation of circuits serving spatial vision and spatial attention (Gattass et al., 1997). The ventral visual-to-hippocampal stream is important for visual memory (Bussey and Saksida, 2007). Multiple lines of evidence have suggested that V2 plays an important role in object recognition memory (Lopez-Aranda et al., 2009). In the monkey, V2 neurons have V1-like (typical Gabor-shaped filter) receptive fields (Liu et al., 2016). Furthermore, mouse V2 neurons have remarkably fine stimulus selectivity, and the majority of response properties in V2 were not different from those in V1 (Van den Bergh et al., 2010). Among the orientation-selective cells in V1 and V2, V1-V2 cortico-cortical connectivity has two types of orientation networks, a tightly synchronized, orientation-preserving network and a less-synchronized orientation diverse network (Roe and Ts’o, 2015), suggesting that V2 neurons may have two distinct functional populations for feature integration.

In our previous study (Long et al., 2021), we have reported visual grid cells in the V2M area from freely foraging rats. This finding, along with the grid-like firing neurons as well as other spatial firing neurons in the S1 (Long and Zhang, 2021, Long et al., 2021, Long et al., 2020), are the first evidence of grid cells in the sensory cortex outside of the MEC from freely foraging rats. The remapping of V2 grid cells has several distinct characteristics from the traditional MEC grid cells. First, MEC grid cells remap in the presence of large changes of the environment (such as a novel room) by causing both translation and rotation, while small changes (changing the enclosure but not the room) do not cause remapping (Hafting et al., 2005, Fyhn et al., 2007). In contrast, V2 grid cells have two distinct subpopulations in terms of the remapping property. Second, the MEC grid cell firing is disrupted in darkness when the visual inputs are removed, even when head-direction cell signaling is preserved (Chen et al., 2016). Moreover, the absence of visual input alters movement velocity modulation of theta frequency. In contrast, the grid-like firing patterns of V2 grid cells are stable. The head directional tuning of V2 grid cells does not change in darkness (**Fig. 2c**).

Two fundamental questions remain: how do the grid-like receptive fields emerge in V2 and what are their functions beyond visual processing? A complete answer to these two questions requires systematic circuit dissection of V2 circuitry and its cell types (Niell, 2015), our findings in the current study provide the first necessary step towards that direction. We will speculate some answers and provide some experimentally testable hypotheses below. The V2 spatial modulation may originate from V1, which may encode rich spatial information (Ji and Wilson, 2007, Haggerty and Ji, 2015, Saleem et al., 2018). Additionally, the source of spatial signal may also arise from the projection from the visual association cortex that has dense projections to the prostrhinal cortex (Burwell and Amaral, 1998), which may transform an animal’s immediate sensory perception of its environment into a spatial map (LaChance et al., 2019). Furthermore, in the visual cortex of macaque monkey, there is an occipito-parietal connection to the posterior parietal cortex (PPC) that has a special role in spatial cognition (Husain and Nachev, 2007, Bicanski and Burgess, 2020). In fMRI imaging, grid-like firing patterns have been reported in multiple areas of human cortices outside the traditional hippocampal formation (Constantinescu et al., 2016, Bao et al., 2019, Doeller et al., 2010, Jacobs et al., 2013). Further, grid-like representations can contribute to mental stimulation in the absence of imagined movement (Bellmund et al., 2016). In animal experiments, grid cells have been reported in the S1 from freely foraging rats (Long and Zhang, 2021). Based on these data, it is reasonable to believe that grid-like firing patterns are broadly distributed across the brain. One new theory for the sensory grid cells is to employ grid cell-like mechanisms to navigate in an abstract “concept” space and learn the structure of the world (objects) (Hawkins et al., 2018), in which the abstract concepts are represented via reference frames. According to Hawkin’s theory, grid-like computation can be implemented in cortical columns in the visual (or somatosensory) cortex to track the location of visual (or tactile) features relative to the objects being viewed (or touched). Together, the neocortex represents object compositionality and learns complete models of objects in different modalities. In V2, there is evidence of column-like compartment that divides neurons with similar receptive field properties (Horton and Adams, 2005). Experimentally, it has also been shown that some V1 and V2 neurons fire only if a visual feature is present at object-centric location on object (Zhou et al., 2000, Williford and von der Heydt, 2016). For instance, V2 neurons in behaving macaques have been showed to encode the side to which the border belongs---the so-called “border ownership coding” during visual processing of complex natural scenes (Zhou et al., 2000). This is reminiscent of the allocentric representation of space in the traditional MEC grid cells. Vision supports the brain’s allocentric spatial navigation (Morris, 1981), it is likely the brain uses both visual and spatial cues to construct a “cognitive map” of the world. The grid-like mechanisms of V2 visual cortex may support the hypothesis that allocentric locations may be used as a basis of visuospatial perception. Together with MEC grid cells and S1 grid cells, V2 grid cells can play the role of location-processing units in navigating complex and sensory cue-rich environments. Recently, it has been shown that the role of MEC grid cells is beyond simple mnemonic coding for space, and can encode behaviorally relevant information such as the reward or goal locality (Boccara et al., 2019, Butler et al., 2019). Very likely, the V2 grid cells may also encode additional spatio-visual information with mixed selectivity for performing a complex task.

Sensory (visual, somatosensory and spatial) cues and velocity signals may contribute to path integration by connecting the “locations” in the sensory space by movements. However, sensory cues and self-motion do not necessarily provide a coherent visuo-spatial or spatio-visual representation. In a recent virtual reality study, visual environmental inputs were dissociated from physical motion inputs, MEC grid cells recorded from mice navigating in virtual open areas showed a greater influence of physical motion, whereas hippocampal place cell firing patterns predominantly reflected visual inputs (Chen et al., 2019). However, it remains unknown whether V2 grid-like patterns are disrupted if the downstream V1 input or the hippocampus is inhibited (e.g., by optogenetic inactivation) during natural navigation. Additionally, unlike MEC grid cells (Stensola et al., 2012), the V2 grid cells are sparsely distributed and show no modular structure between the deep and superficial layers. It is also important to note that there are heterogeneous spatial tunings in V2, as seen by simultaneously recorded V2 grid cells, head-direction cells, place cells and border cells.

Remapping is a property of neural plasticity in response to changes in environment and sensory cues. Grid firing patterns of traditional MEC grid cells can remap (such as grid rotation and rescaling) in the response to not only the environmental change (Fyhn et al., 2007), but also the task learning (Butler et al., 2019). In our study, rats are freely foraging in the environmental enclosure without performing an explicit task. The remapping of bistable V2 grid fields predominantly preserved their firing rates, gridness scores and grid radius, but changed the fields by rotation and translation. Conjunctive head-direction tuning of bistable V2 grid cells was also remapped (**Fig. 2**). Notably, the bistable remapping in V2 grid cells has not been found in other simultaneously recorded V2 spatially tuned cells, neither in the MEC grid cells or S1 grid cells. Therefore, the functional significance of the bistable remapping mechanism of V2 grid cells remains mysterious, as we did not find any significant differences in grid cell characteristics (such as the gridness score, grid radius, spatial frequency, and theta modulation index) between the V2 bimodal and regular grid cell populations. However, we cannot exclude the possibility this may be a sampling issue.

To understand the computational mechanism of neuronal representations, grid cell’s receptive field can be closely approximated by a two-dimensional cosine function with a specific spatial frequency. Therefore, grid cells can perform discrete Fourier transform-like computation (Kubie and Fox, 2015, Rodriguez-Dominguez and Caplan, 2019). Changes by rotation and translation may be interpreted as an angular modulation in the frequency domain. Several computational models and theories have been proposed for the MEC grid cells and remapping (Hardcastle et al., 2017, Stachenfeld et al., 2017, Mosheiff and Burak, 2019, Agmon and Burak, 2020, Anselmi et al., 2020, D’Albis and Kempter, 2020); for a review, see (Giocomo et al., 2011, Zilli, 2012). The “oscillatory interference” model explains the theta phase precession (Burgess et al., 2007), whereas the continuous attractor models can generate grid firing patterns (Burak and Fiete, 2009, Bush and Burgess, 2014). The underlying computational model explaining the remapping bistability for V2 grid cells needs further exploration.

## Supplemental information

Supplemental information includes ten figures.

## Acknowledgments

We thank Edvard Moser, May-Britt Moser, Daniel Bush and Neil Burgess for sharing their analytical codes with us. S.-J.Z. is supported by the National Natural Science Foundation of China (Grant# 31872775). Z.S.C. is partly supported by the US National Institutes of Health (R01- MH118928).

## Author contributions

S.-J.Z. conceived the project. X.L. and S.-J.Z. designed the study, performed the surgery and analyzed the data. J.C., B.D. and X.L. conducted the experiments and acquired the recordings. B.D. and Z.S.C. provided the feedback. X.L. developed the codes and made the figures. Z.S.C., X.L. and S.-J.Z. wrote the manuscript.

## MATERIALS AND METHODS

### Experimental subjects

Eight experimentally naïve male adult Long-Evans rats (2 to 4 months old, 250 to 450 grams at the time of implantation, six rats for V2 implanting and two rats for MEC implanting) were used in this study. Prior to surgery, rats were group-housed with four male littermates per cage. After chronic implantation, animals were housed singly in transparent cages (35 cm x 45 cm x 45 cm, W x L x H) and kept on a 12- hour light/12-hour dark cycle (lights on at 9 p.m. and off at 9 a.m.). Rats were maintained in a temperature- (19-22°C), humidity- (55-65%) controlled vivarium. One week after postoperative recovery, animals were food-deprived 8-24 hours before behavioral training or experimental recording trials and were kept at ∼90% of free-feeding body weight with free access to water. Experimental training and recording were performed during the dark phase of the cycle. All animal experiments were conducted according to the pre-approved protocol by the Animal Care and Use Committee from both Army Medical University and Xinqiao Hospital in accordance with the National Animal Welfare Act.

### Surgery and electrodes

Rats were given intramuscular injection of buprenorphine (0.001 mg per kg) just right before chronic surgery. Animals were anaesthetized under 1.5-3% of isoflurane in O2, immobilized in a stereotaxic frame (David Kopf Instruments, California, USA) and maintained on a feedback-controlled heating plate at 37°C. Four tetrode bundles based on 16-channel microdrives (Axona, St. Albans, UK) were constructed with 17 µm Platinum/Iridium wires (#100167, California Fine Wire Company, USA). The electrode tips were cut and platinum-electroplated to reduce their target impedance to 150 to 300 kΩ measured at 1 kHz using an electroplating plater (nanoZ; White Matter LLC, Seattle, Washington, USA). Microdrives were implanted to target the mediolateral (V2ML) and mediomedial (V2MM) regions of the second visual cortex (V2) with the stereotaxic coordinate centered at ∼2.5 mm lateral to the midline (ML), ∼−4.5 mm anterior-posterior (AP) from bregma, 0.4-2.0 mm dorsal-ventral (DV) below the dura and at an angle of 10° from the medial-to-lateral direction in the coronal plane. Implanted microdrives were anchored to jeweler’s screws in the skull and secured with multiple rounds of application of dental cement. The electrode was grounded via a reference electrode connected with one jeweler’s screw fixed to the skull.

### Behavioral and recording procedures

Animals were allowed at least one week for postsurgical recovery in their home cages. During the behavioral training, rats were handled three times a day to get familiar with the testing arena and experimenter. Animals were trained to run in different shapes or sizes of open field enclosures. Implanted animals were individually transferred to the experimental recording room with a covered carrying container. Rats rested in the flowerpot on a holding platform outside the recording box before and after each recording session. Rats were mildly disoriented and then placed back to the open filed enclosure box from a random entry direction during each recording trial. Microdrives mounted on animals’ heads were connected to the recording device (Axona Ltd, St Albans, UK) and equipped with a headstage and 5-m lightweight cable. A white cue card (297 mm x 210 mm) was used as a single polarizing cue and taped to the interior enclosure wall. Each daily recording session lasted at least 20 min to facilitate full coverage of the testing enclosure while they actively foraged for food pellets scattered into the enclosure. Tetrodes were advanced towards deep layers of the second visual cortex in rats during each recording session. Electrodes were lowered in steps of 25 or 50 µm per day until well-isolated single units can be identified. Unit signals were acquired by the DacqUSB system (Axona Ltd., St. Albans, U.K.) digitized at 48 kHz, band-pass filtered between 800 Hz and 6.7 kHz and a gain of x5-18k. Local field potentials were recorded from one of the electrodes with a low-pass filter (500 Hz).

### Spike sorting and cell classification

Spike sorting was manually performed offline with graphical cluster-cutting software TINT (Neil Burgess and Axona Ltd, St. Albans, U.K.), and the clustering was primarily based on features of the spike waveform (peak-to-trough amplitude and spike width), together with additional autocorrelations and cross-correlation. To make sure that the same neuron was not counted twice, signal units with similar or identical waveform shapes were counted only once during our manual cluster cutting, whenever similar or identical individual neurons were recorded and tracked across successive recording sessions. To confirm the quality of cluster separation, we calculated L-ratio as well as isolation distance between clusters. Two small infrared light-emitting diodes (LEDs) were attached to the rat’s head to track the rats’ speed, head position and orientation using an overhead video camera and tracking hardware/software. Only spikes with instantaneous running speeds > 2.5 cm/s were excluded for further analysis in order to exclude confounding behavioral effects such as immobility, grooming and rearing. To construct firing fields and firing rate distributions, the position data were divided into 2.5-cm x 2.5-cm location bins, and the path was smoothed with a 21-sample boxcar window filter (400 ms; 10 samples on each side).

Cells with > 100 spikes per session and with a coverage of > 80% were included for further analyses. Maps for the number of spikes and dwell time were smoothed with a quasi-Gaussian kernel over the surrounding 5 x 5 bins. Spatial firing rates were calculated by dividing the number of spikes with dwell time. The peak firing rate was constructed as the highest rate in the corresponding bin from the spatial firing rate map. Mean firing rates were measured from the whole recording session data.

### Analysis of place cells

Spatial information is a quantification used to determine the extent to which a neuron’s spatial firing pattern can predict the actual position of freely moving animals and is expressed in bits/spike. The spatial information was calculated as:

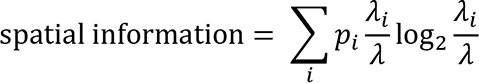

where *λ*_*i*_ is the mean firing rate of the given cell in the *i-*th bin, *λ* is the overall mean firing rate of the cell in the whole recording trial, and *p*_*i*_ is the probability for the rat being at the spatial location of the

*i-*th bin.

Adaptive smoothing (Skaggs et al., 1996) was used to optimize the trade-off between sampling error and spatial resolution before the calculation of spatial information. The position data were first sorted into 2.5-cm x 2.5-cm bins, and to calculate the firing rate for a specific bin, circle centered on the given bin is expanded gradually until

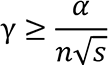

where γ is the circle’s radius in bins, *n* and *s* are the total number of occupancy samples and spikes within the circle, respectively, and α =10,000. With a video sampling frequency at 50 Hz, the firing rate of the bin was then set to 50 · *s*/*n*. A place cell was defined with its spatial information above the chance level, which was computed by a random shuffling procedure using the total sample of recorded cells. For each round of the permutation process, the entire sequence of spike trains was time-shifted along the animal’s trajectory by a random amount of time between 20 s and the total trial duration minus 20 s, with the end of the spike train wrapped to the beginning of the trial. A spatial firing map was then constructed, and for each shuffled map, spatial information was calculated. This permutation process was repeated 100 times for each cell, leading to a total of 136,400 permutations for the 1364 visual neurons. This shuffling procedure aimed to preserve the temporal firing patterns while perturbing the spatial structure at the same time.

The threshold for classifying cells into place cells was defined as the spatial information scores exceeding the 99^th^ percentile of the entire shuffled data.

### Analysis of grid cells

Visual grid cells were identified by calculating the 6-fold symmetry in their spatial autocorrelations with smoothed rate maps (Boccara et al., 2010, Sargolini et al., 2006, Zhang et al., 2013).

With *λ*(*x*, *y*) representing the mean firing rate of a given cell at coordinate (*x*, *y*), the autocorrelation between the firing field itself and the firing field with spatial lags of *λ*_*x*_ and *λ*_*y*_ was determined as:

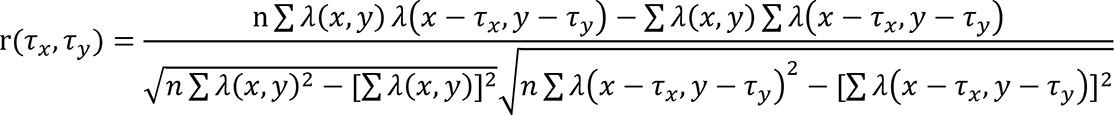

where the summation is over total n pixels in *λ*(*x*, *y*) for which firing rate was estimated for both *λ*(*x*, *y*) and *λ*(*x* − *λ*_*x*_, *y* − *λ*_*y*_). Autocorrelations were not calculated for spatial offsets when total number of overlapping pixels was smaller than 20. The degree of spatial regularity (“gridness” or “grid score”) was estimated for every single unit by using an annular sample surrounding the central peak of the autocorrelogram but excluding the central peak itself, and by comparing rotated versions of this annular sample (Boccara et al., 2010, Sargolini et al., 2006). The Pearson’s correlations between this annular sample and its rotated versions were obtained, with the angles of shift being 60° and 120° in the first group, and rotation degree of 30°, 90° and 150° in the second group. Gridness or the neuron’s grid score was characterized as the minimal difference between any of the correlation coefficients in the first rotation group and any of the correlation coefficients in the rotation second group. Shuffling was performed in the same way used for defining place cells. For each permutation, a randomized grid score was obtained. Grid cells were categorized as cells with grid scores exceeding the 99^th^ percentile of all randomized grid scores from shuffled data over the entire population of identified visual cells. Grid spacing was defined as the median distance from the central peak of the autocorrlogram to the closest vertices among six adjacent spatial firing fields in the autocorrelogram of the spatial firing map. Since such analysis is rather sensitive to noise in the autocorrelogram, grid spacing was determined only when the median distance to the six surrounding vertices for the cell was comparable to the grid radius, which was defined as the radius of the circle resulting in the highest grid score. Thus, the grid radius was also defined as the grid field size.

The orientation of the gird cell was calculated by first defining the camera-based horizontal reference line of being 0°. The vector from the central peak to the nearest surrounding vertices of the autocorrelogram in the counterclockwise direction was defined as grid orientation.

### Analysis of head-direction cells

The rat’s head direction was quantified by the relative position of the two diodes differentiated through their sizes (Boccara et al., 2010, Sargolini et al., 2006, Zhang et al., 2013). The directional tuning curve for a given cell was drawn by calculating the firing rate of the cell as a function of the rat’s heading direction, which is divided into angular bins of 3 degrees and subsequently smoothed with a 15-degree mean window filter. To avoid possible bias, recorded data were only used if all head directional bins contain spikes.

The strength of directionality was estimated by computing the mean vector length from the polar plot of angular firing rates. The chance values or the threshold for defining head-direction cells were determined by a shuffling procedure in a similar way as for place cells, with the entire sequence of spike trains shifted between 20 s and the total trail length minus 20 s along the animal’s path. Cells were classified as head-direction cells if their mean vector lengths surpass the 99^th^ percentile of the mean vector lengths of the entire shuffled distribution.

### Analysis of border cells

Putative border or boundary vector cells were identified by computing the border score, which is defined as the difference between the maximum coverage of a given wall by any single spatial firing field of the recorded cell and the average firing distance of all the pixels of the firing field to the nearest wall, normalized by the sum of those two values (Boccara et al., 2010, Zhang et al., 2013, Solstad et al., 2008). Border scores ranged from −1∼+1, with -1 representing perfect central firing fields and +1 representing spatial firing fields exactly lining up with at least one entire wall. Firing fields were defined as total number of neighboring pixels with firing rates exceeding 0.3 times the highest firing rate that covered a total area more than 200 cm^2^.

The definition of border cell was in a similar way as for place cells, head-direction cells and grid cells. For each permutation process, the whole sequence of spike trains was shifted along the animal’s path by a random interval between a period of 20 s and the total time of the entire trial minus 20 s, with the end of the spike train wrapped to the start of the trial. A spatial firing rate map was then derived, and a border score was estimated.

The distribution of border scores was obtained for the entire set of shuffling trials from the whole population, and the 99^th^ percentile was then determined as the classifying threshold. Border cells were defined with observed border score higher than the threshold value.

### Histology and identification of tetrode trajectory

At the completion of the last recording session for each implanted animal, rats were deeply anesthetized by giving an overdose of sodium pentobarbital. Rats underwent transcardial perfusion first with phosphate-buffered saline (PBS) followed by 4% paraformaldehyde (PFA). Brains were extracted and post-fixed in 4% PFA overnight. The individual brain was then transferred in 10, 20 and 30% sucrose/PFA solution sequentially across 72 hours. After brains sank, they were sectioned at 30 μm intervals through the tetrode implant region with a cyrotome. Coronal sections were mounted on gelatin-coated glass slides and stained with Cresyl violet (Sigma-Aldrich). The final tetrode trajectories through V2 subregion were identified through 3D reconstruction of digitized images of the Cresyl violet-stained coronal sections acquired with the Olympus Slideview VS200 Digital Slide Scanner. Tetrode trajectories of each individual recordings were measured from the deepest tetrode location, daily advancement for tetrode lowering and adjusted tissue shrinkage correction by dividing the distance between the initial brain surface and the final electrode tip by the last advanced depth of the recording electrodes. Electrode traces were confirmed to be located within the secondary visual cortex in rats according to The Rat Brain in Stereotaxic Coordinates (Paxinos and Watson, 2013).

---

**Fig. S1.**
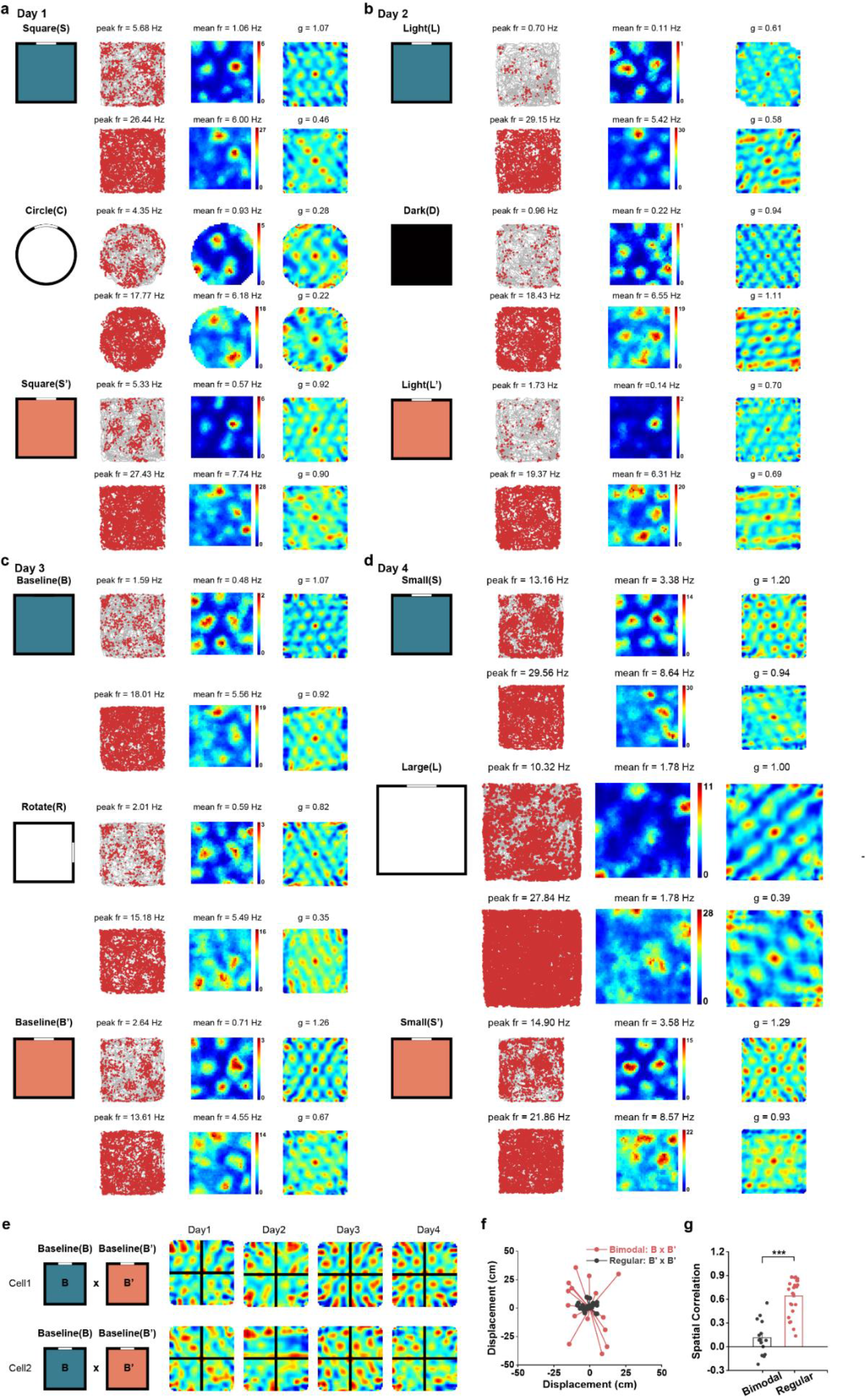
V2 regular grid cells did not switch firing patterns during change in the familiar environment. Spatial responses of co-recorded V2 regular grid cells. (a) Grid firing patterns in different shapes of running boxes. (b) Grid firing patterns maintained under total darkness. (c) Grid firing patterns were consistent after cue card manipulation. (d) Grid firing patterns remain unaltered in different sizes of running boxes. (e) Cross-correlogram for the same cell in the same baseline condition after environmental change (B versus B’). Note the peak of the cross-correlogram was nearly centered on the origin. (f) Vector diagram showing the grid displacement from the center of the cross-correlogram to the nearest peak. The mean displacement for regular grid cells was significantly higher than that for bimodal grid cells. (g) Comparison of spatial correlation coefficients between two baselines after environmental manipulation (B versus B’), with significantly higher correlation for regular grid cells than bimodal grid cells.

**Fig. S2.**
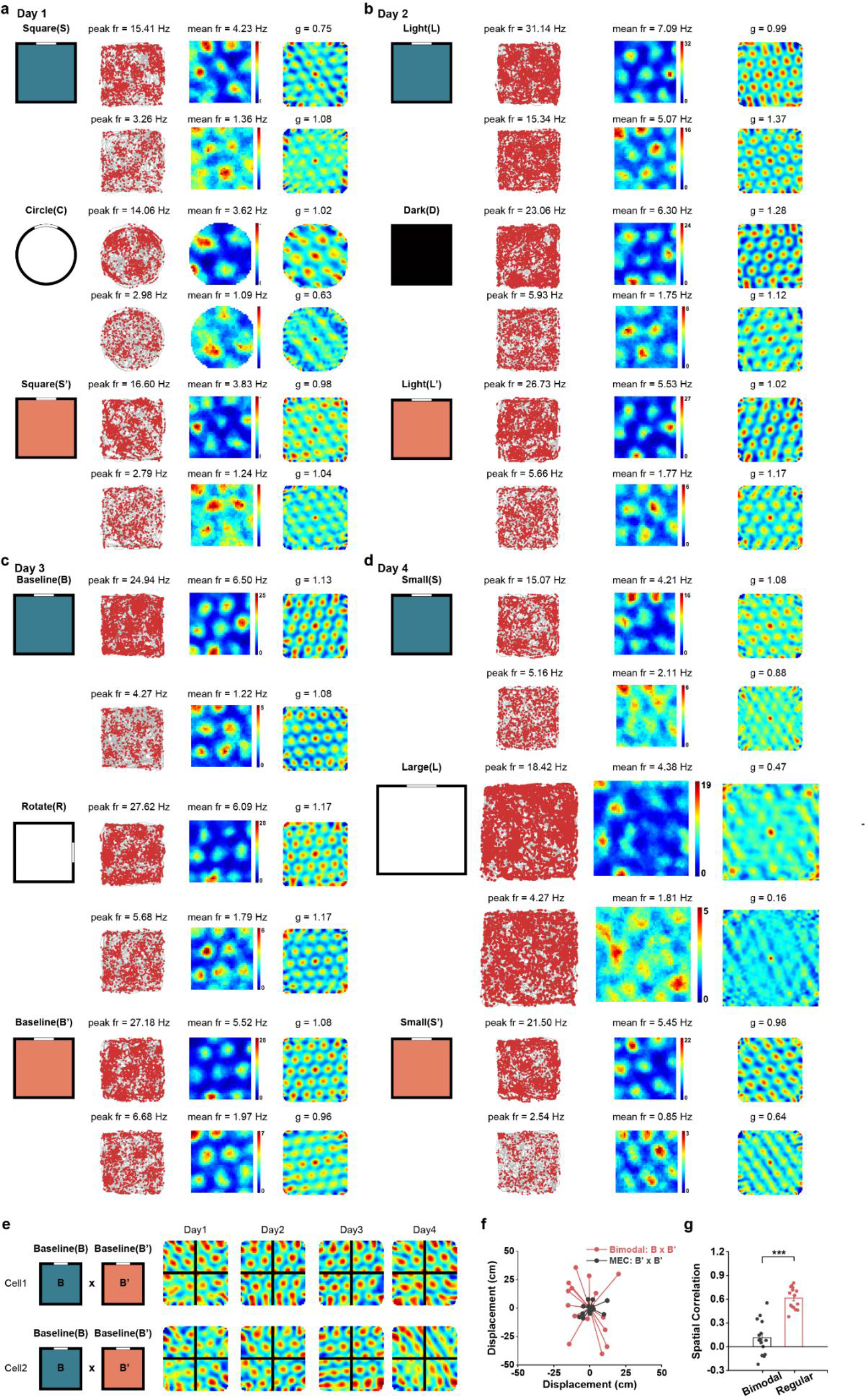
MEC grid cells did not switch firing patterns during two baseline sessions after change in the familiar environment. Spatial responses of co-recorded one pair of MEC grid cell. (a) Grid firing patterns in different shapes of running boxes. (b) Grid firing patterns maintained under total darkness. (c) Grid firing patterns were consistent after cue card manipulation. (d) Grid firing patterns remain unaltered in different sizes of running boxes. (e) Cross-correlogram for the same cell in the same baseline condition after environmental change (B versus B’). Note the peak of the cross-correlogram was nearly centered on the origin. (f) Vector diagram showing the grid displacement from the center of the cross-correlogram to the nearest peak. The mean displacement for regular grid cells was significantly higher than that for bimodal grid cells. (g) Comparison of spatial correlation coefficients between two baselines after environmental manipulation (B versus B’), with significantly higher correlation for MEC grid cells than V2 bimodal grid cells.

**Fig. S3.**
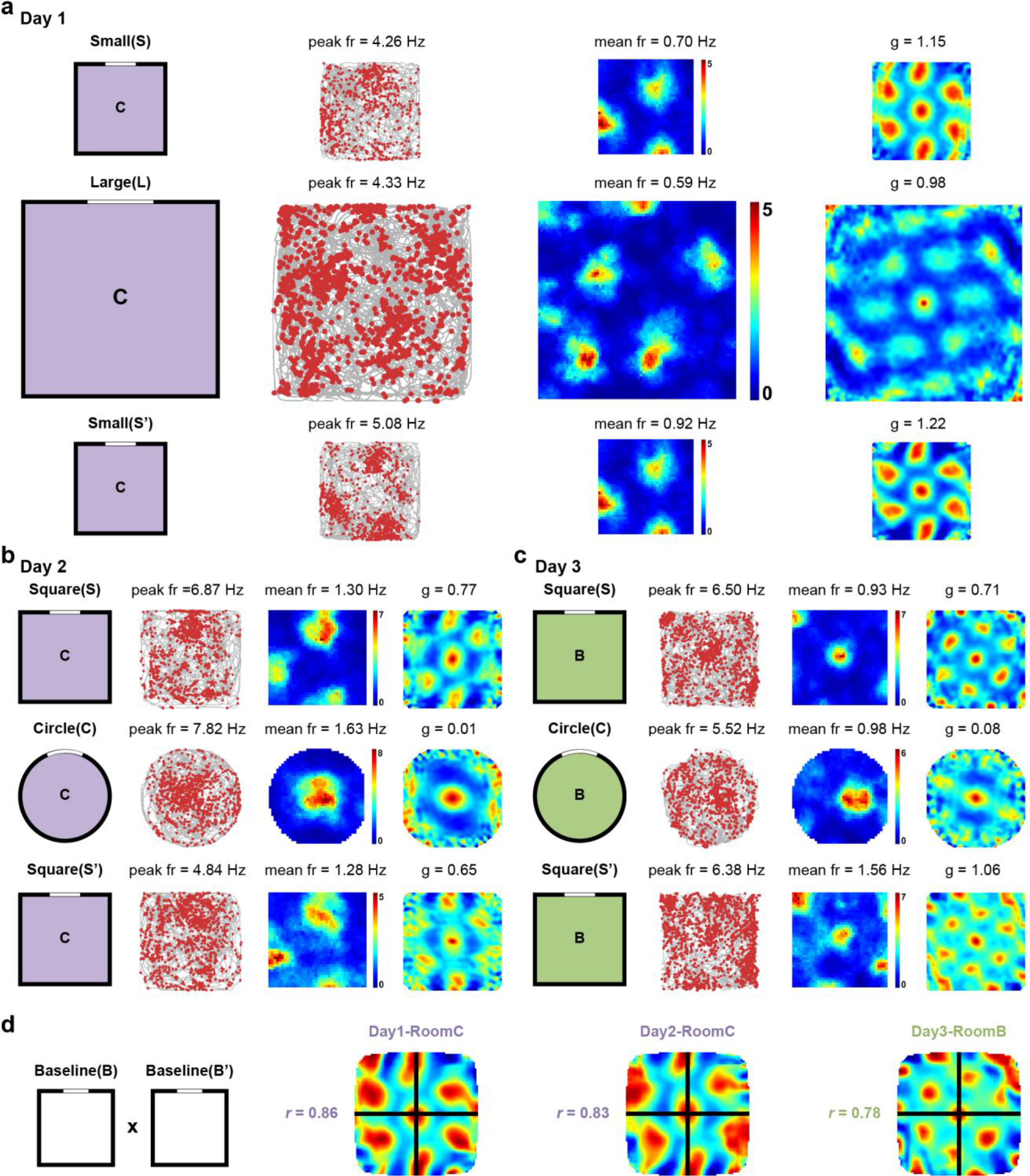
Bimodal grid cells did not switch firing patterns in the novel environment. Spatial responses of the same cell as in **Figure 1** except in two novel environments. (a) Grid firing patterns in different shapes of running boxes in novel room C. (b) Grid firing patterns remain unaltered in different sizes of running boxes in novel room C. (c) Grid firing patterns did not switch in different shapes of running boxes in novel room B. (d) Cross-correlogram for the same cell in the same baseline condition after environmental change (B versus B’). Pearson correlation coefficients (*r*) of the cross-correlation between spatial maps of the two baseline sessions are indicated. Note the peak of the cross-correlogram was nearly centered on the origin.

**Fig. S4.**
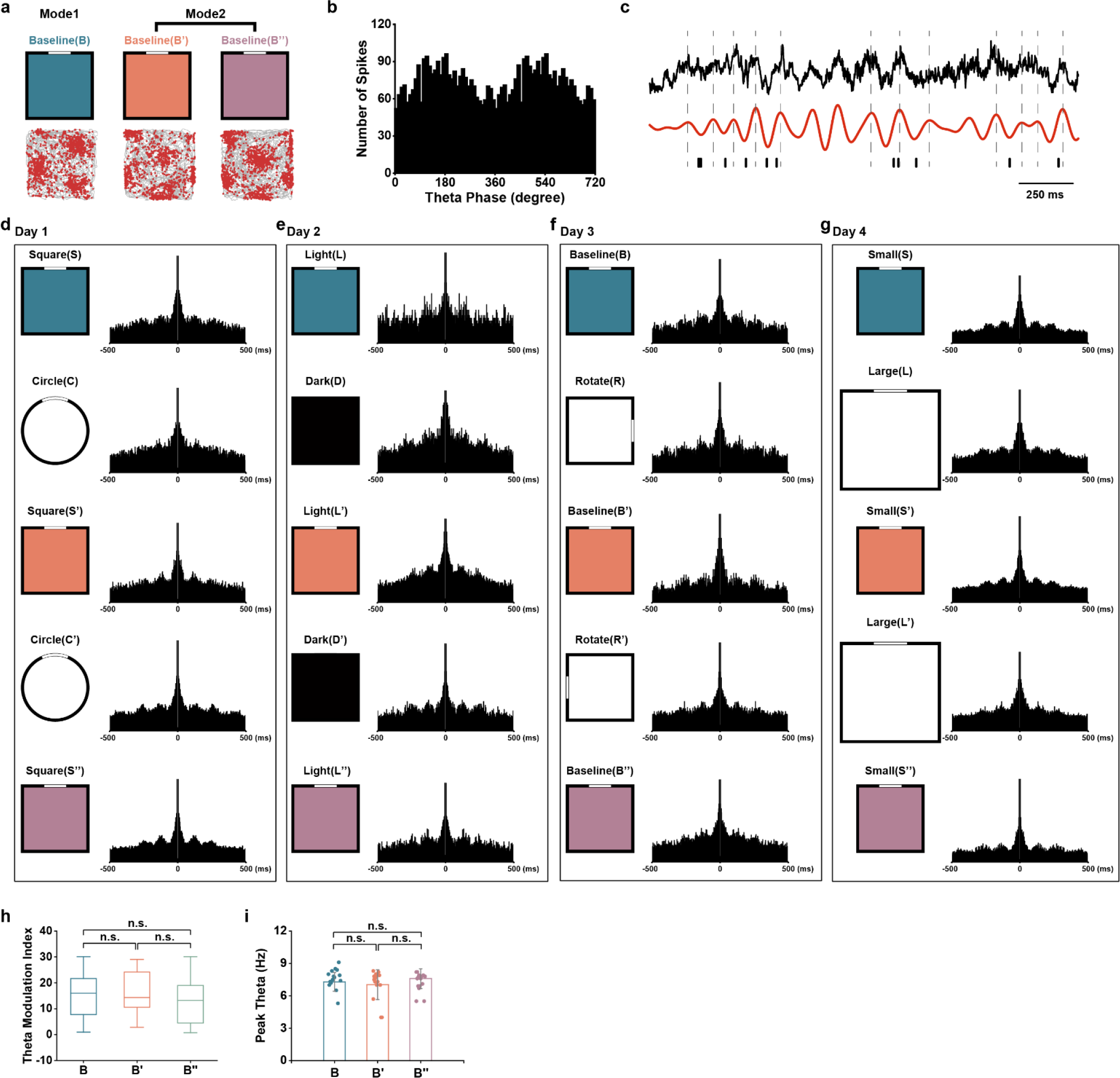
Theta modulation preserved during bimodal switch of firing patterns of grid cells by environmental change. (a) Bimodal firing fields of the same grid cell in **Fig. 1**. (b) Distribution of firing rate within two theta cycles (peak of local theta rhythm set as 0°). (c) LFP signals with spike times for the grid cell during 2 s of active running. Top, unfiltered EEG trace; middle, theta (6-12 Hz) filtered trace; bottom, individual spikes. Gray dashed lines indicate 0° of theta phase. (**d**-**g**) Theta modulation of the same grid cell as in **Fig. 1** during bimodal switch of spatial firing patterns in responses to different environmental changes. (h) Comparison of theta modulation index in the baseline condition (B), after environmental manipulation and back to the baseline condition (B’) and after another environmental change and back to the baseline condition again (B’’). n.s., not significant. (i) Same as (**h**) except for theta peak frequency.

**Fig. S5.**
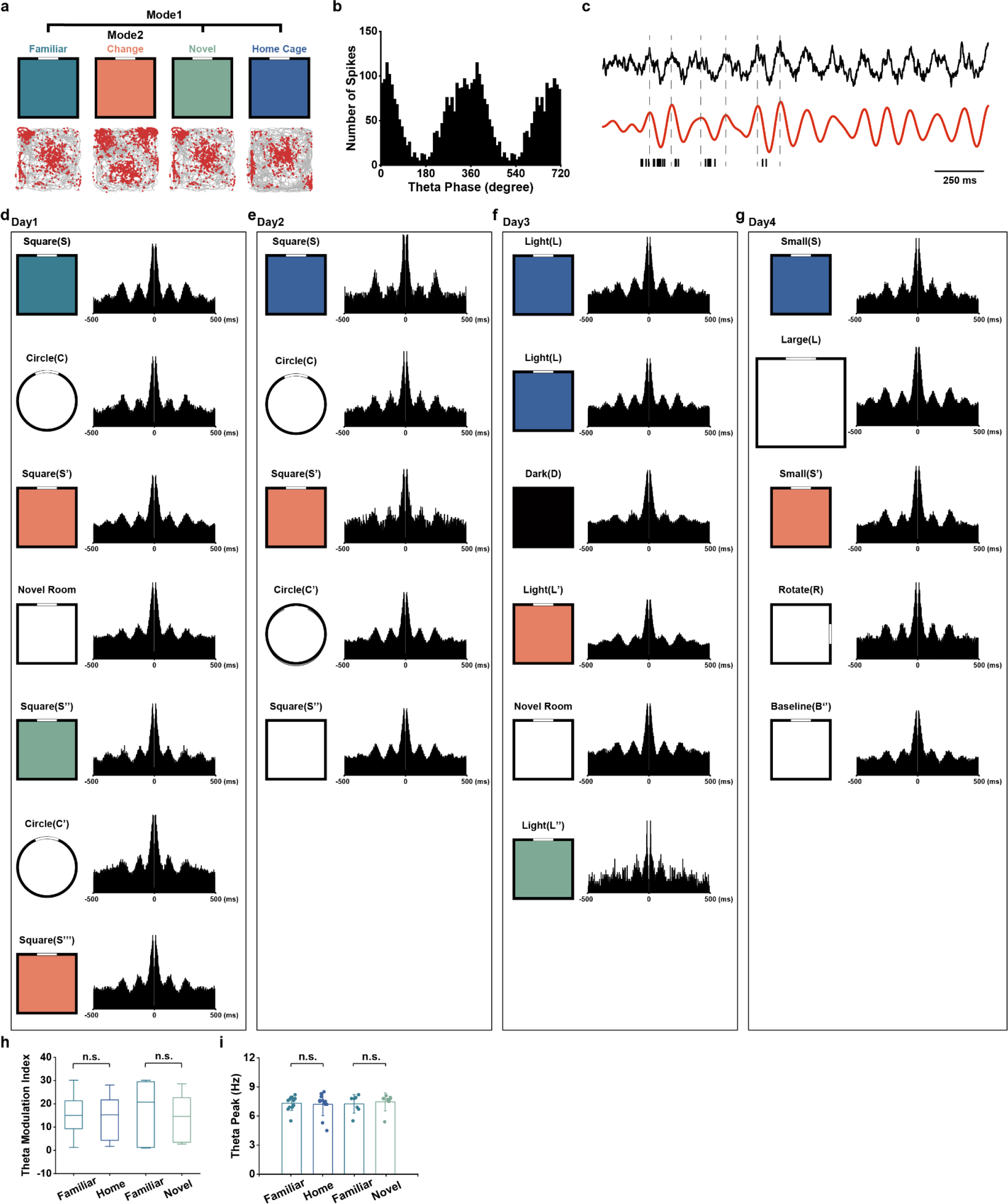
Theta modulation persisted during re-entry into the same familiar environment. (a) Bimodal firing fields of the same grid cell in **Fig. 2**. (b) Distribution of gird firing rate within two theta cycles (peak of local theta rhythm set as 0°). (c) LFP signals with spike times for the grid cell during 2 s of active running. Top, unfiltered EEG trace; middle, theta (6-12 Hz) filtered trace; bottom, individual spikes. Gray dashed lines indicate 0° of theta phase. (**d**-**g**) Theta modulation and theta phase distribution of the same grid cell as in Figure. 2 during bimodal switch of spatial firing patterns in responses to different environmental changes and re-entry into the same familiar environment either from the home cage or from a novel room. (h) Comparison of theta modulation index in the baseline condition (B), after environmental manipulation and back to the baseline condition (B’) and after another environmental change and back to the baseline condition again (B’’). n.s., not significant. (i) ) Same as (**h**) except for theta peak frequency.

**Fig. S6.**
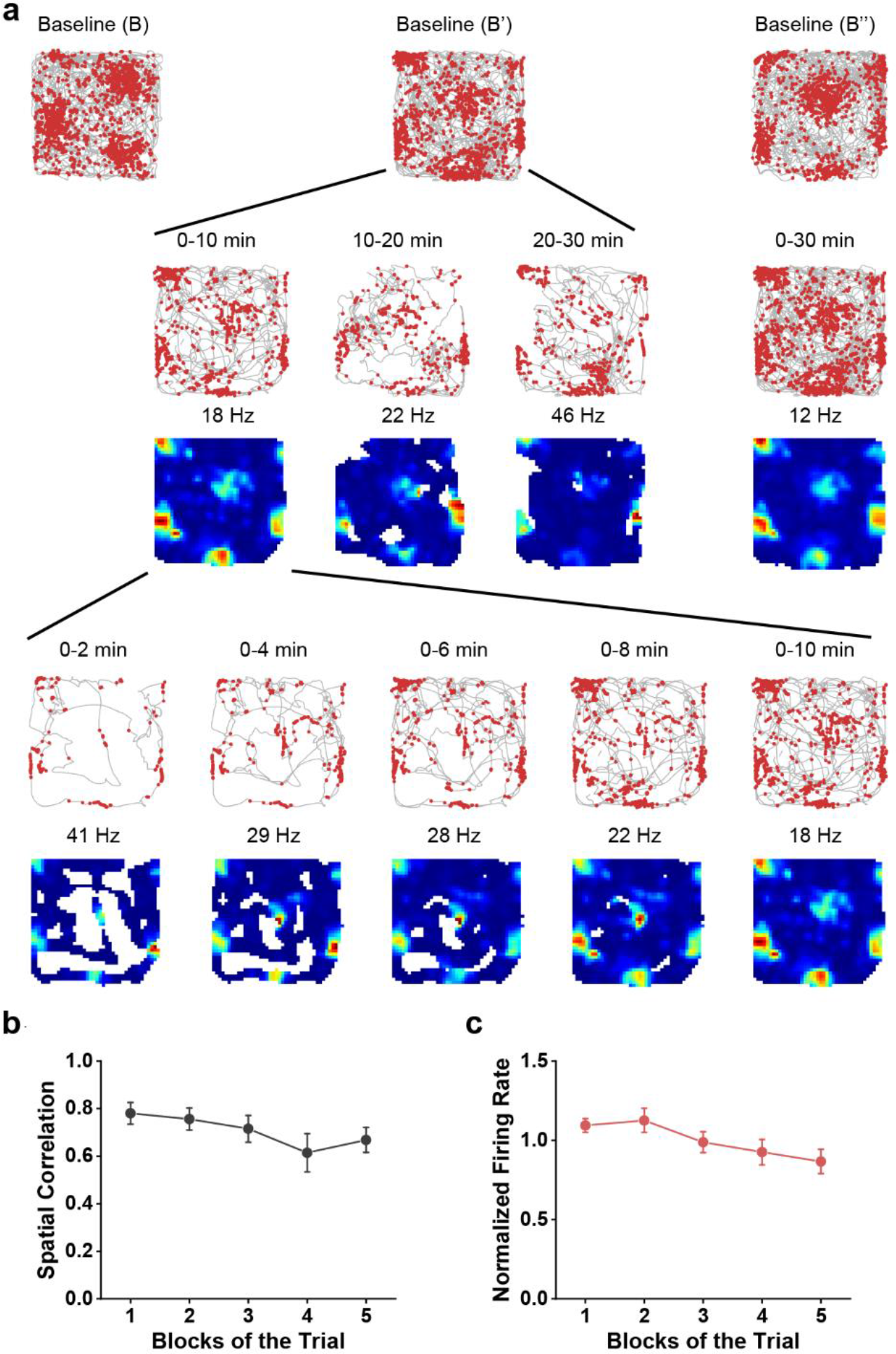
Bimodal shift of grid firing pattern is expressed instantly after environmental change. (**a**), Trajectory maps for the same grid cell in Fig. 1a in a baseline condition (B), after environmental manipulation and back to the baseline condition (B’) and after another environmental change and back to the baseline condition again (B’’). The middle trial of the baseline condition (B’) is broken into blocks to show the development of grid structure after bimodal shift. (b) Development of grid firing patterns as shown by spatial correlation with the whole session of the firing field of the baseline trial (B’) across preceding 5 divided blocks of the same trial (*n* = 15, means ± s.e.m.). (c) Stability of mean firing rate across 5 divided blocks, normalized to the mean firing rate for the whole session.

**Fig. S7.**
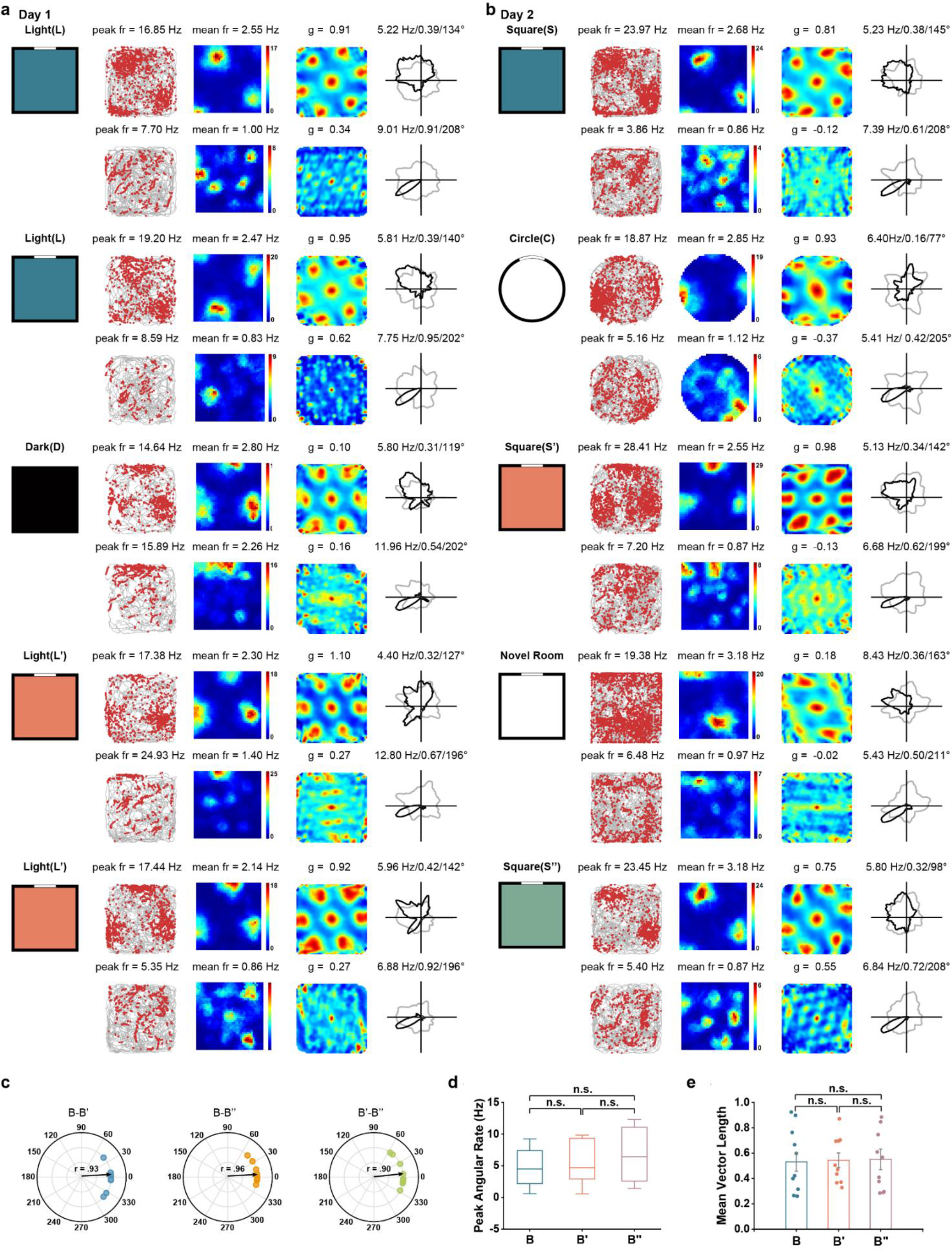
Head directionality preserved during bimodal shift of grid firing patterns. (**a**-**b**) Co-recorded representative conjunctive grid cell and head-direction cell during bimodal switch of grid firing patterns. The head directionality of one conjunctive V2 grid cell and one V2 head-direction cell preserved while the grid pattern switched between two different spatial firing patterns. (**c**) Polar plots showing offset of peak angular direction of conjunctive grid cells (*n* = 4) and head-direction cells (*n* = 6) during bimodal switch of grid firing patterns. (**d**-**e**) The peak angular rate and mean vector length showed no significant change during bimodal remapping of V2 grid cell firing patterns.

**Fig. S8.**
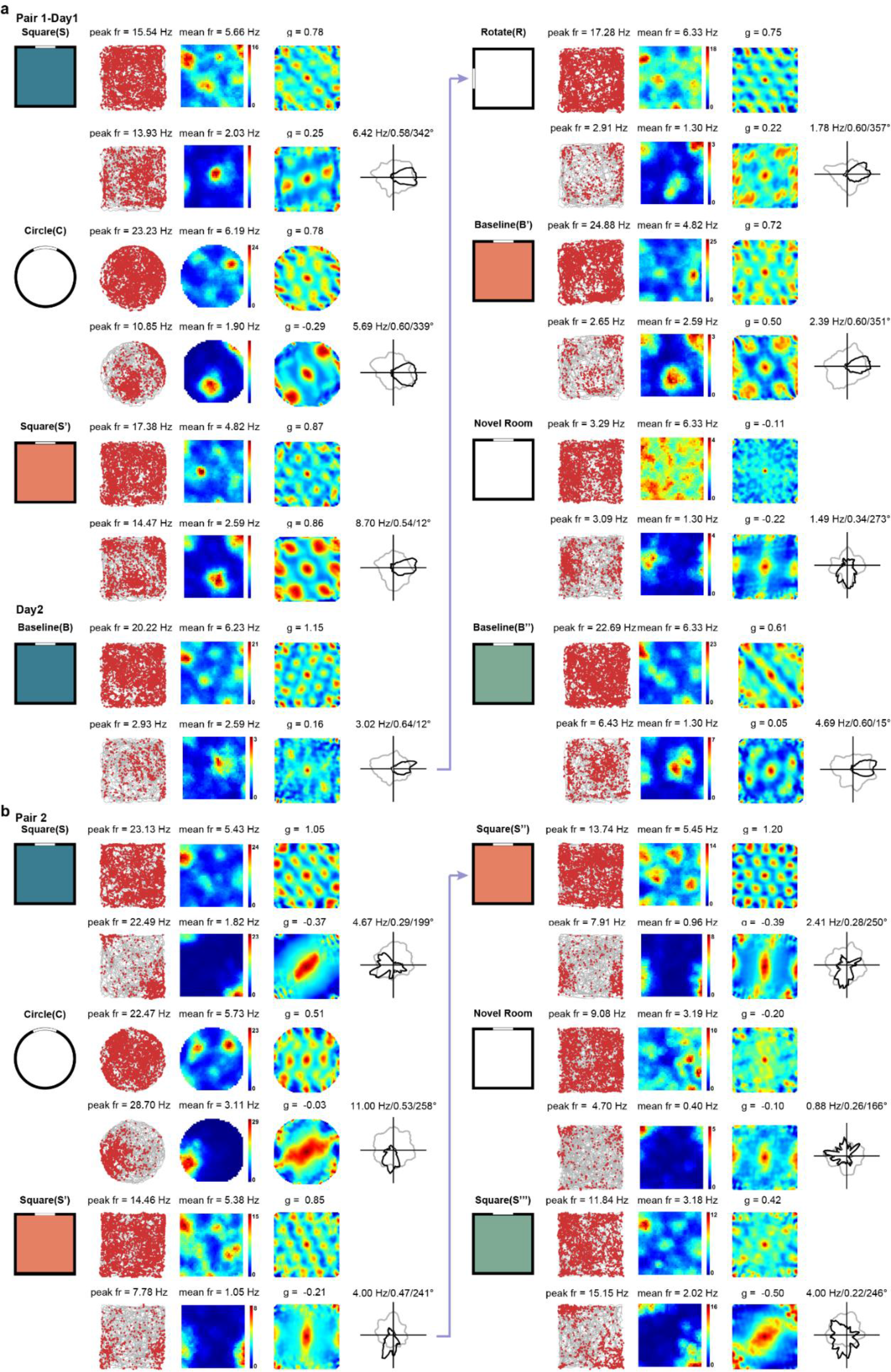
Co-vary and co-restoration of grid firing patterns of two pairs of co-recorded V2 bimodal grid cells. **(a)** Simultaneously recording of a bimodal grid cell and a conjunctive bimodal grid cell switch their spatial firing pattern when the familiar environment change, and restore the same firing pattern when re-entry into the same familiar environment from a novel room. Note that the head directionality of the conjunctive bimodal grid cell remained unchanged despite the bimodal switch of spatial firing pattern. **(b)** Simultaneously recorded a bimodal grid cell and a conjunctive bimodal grid cell switch their spatial firing pattern when the familiar environment changes.

**Fig. S9.**
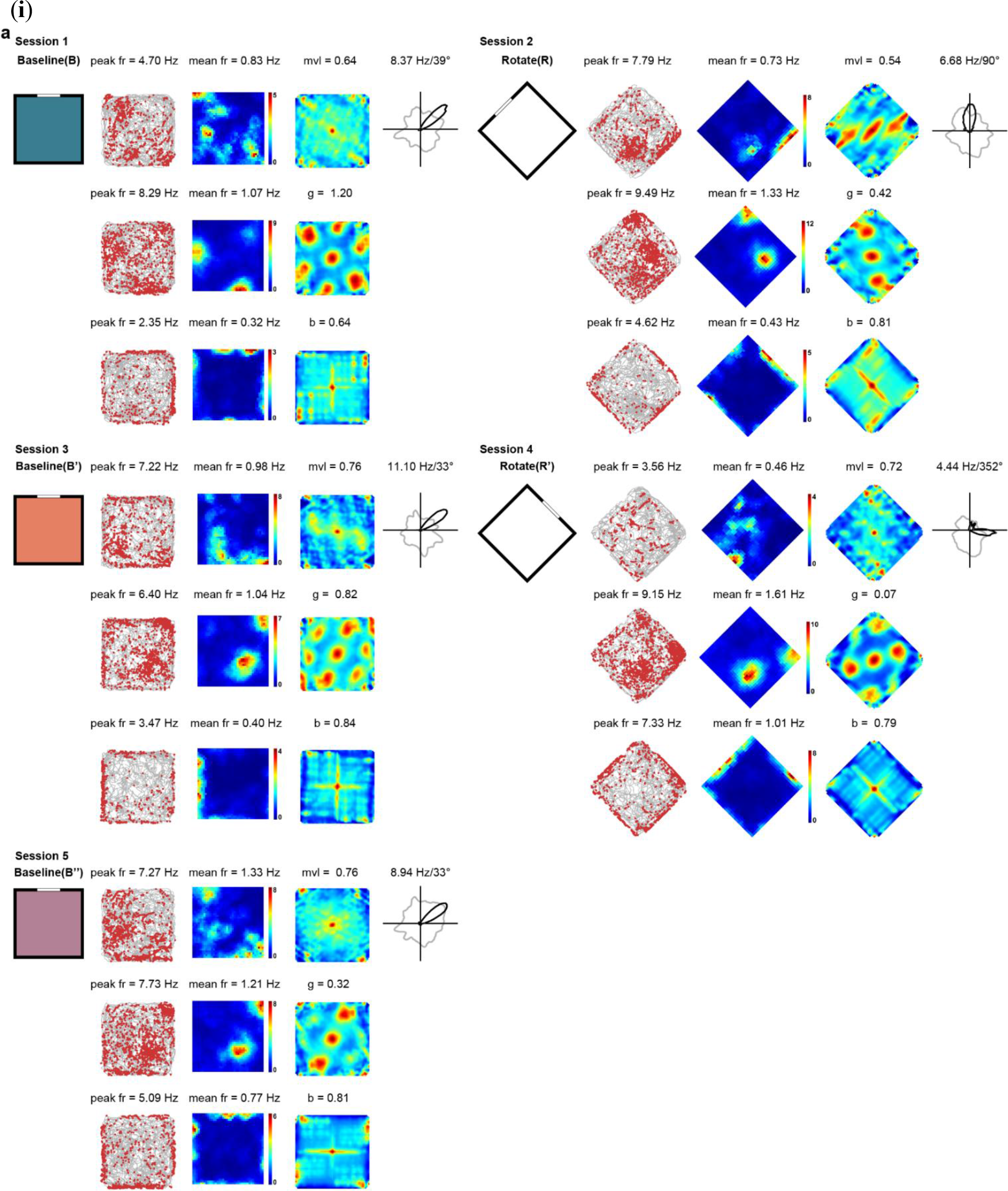
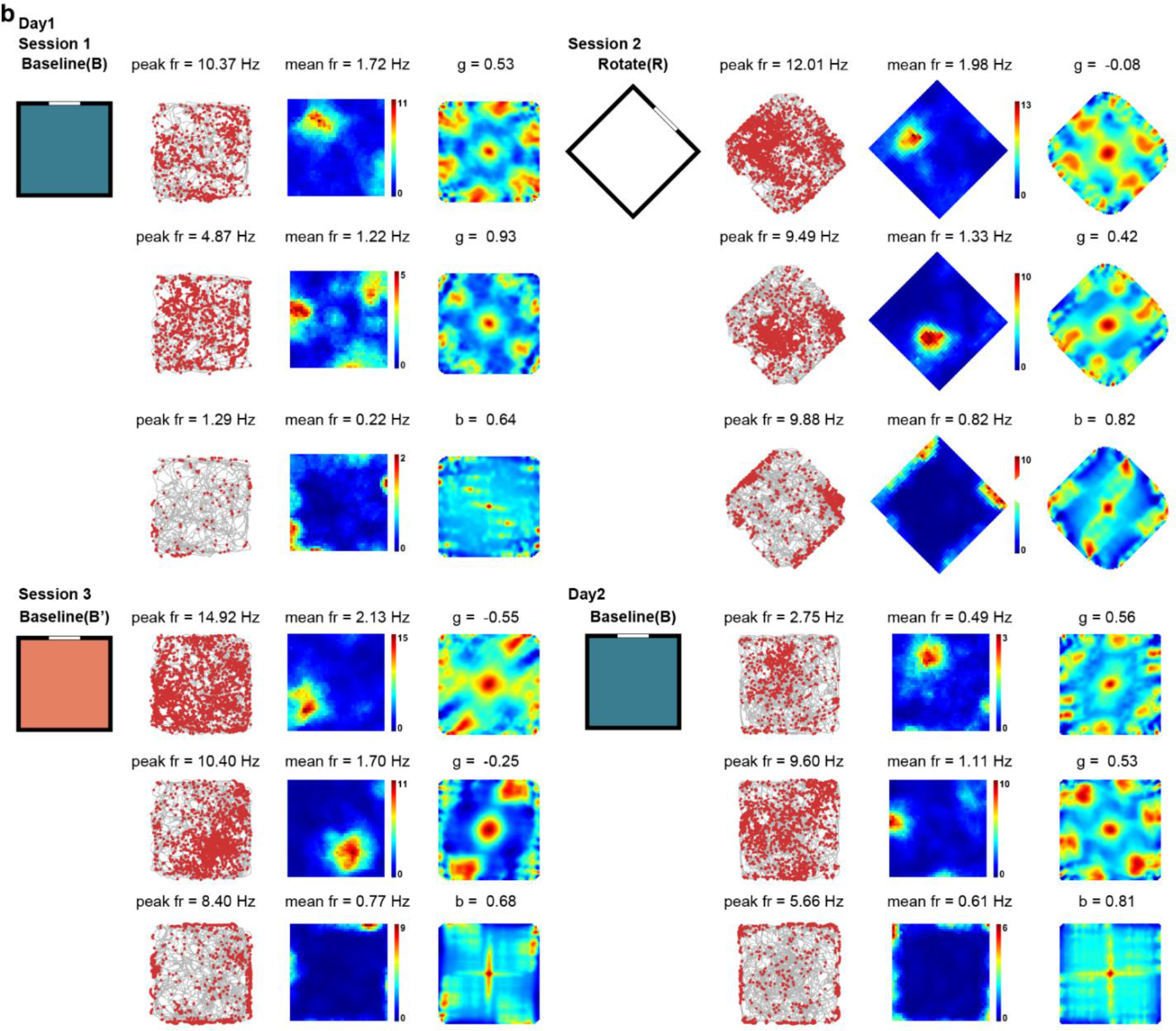
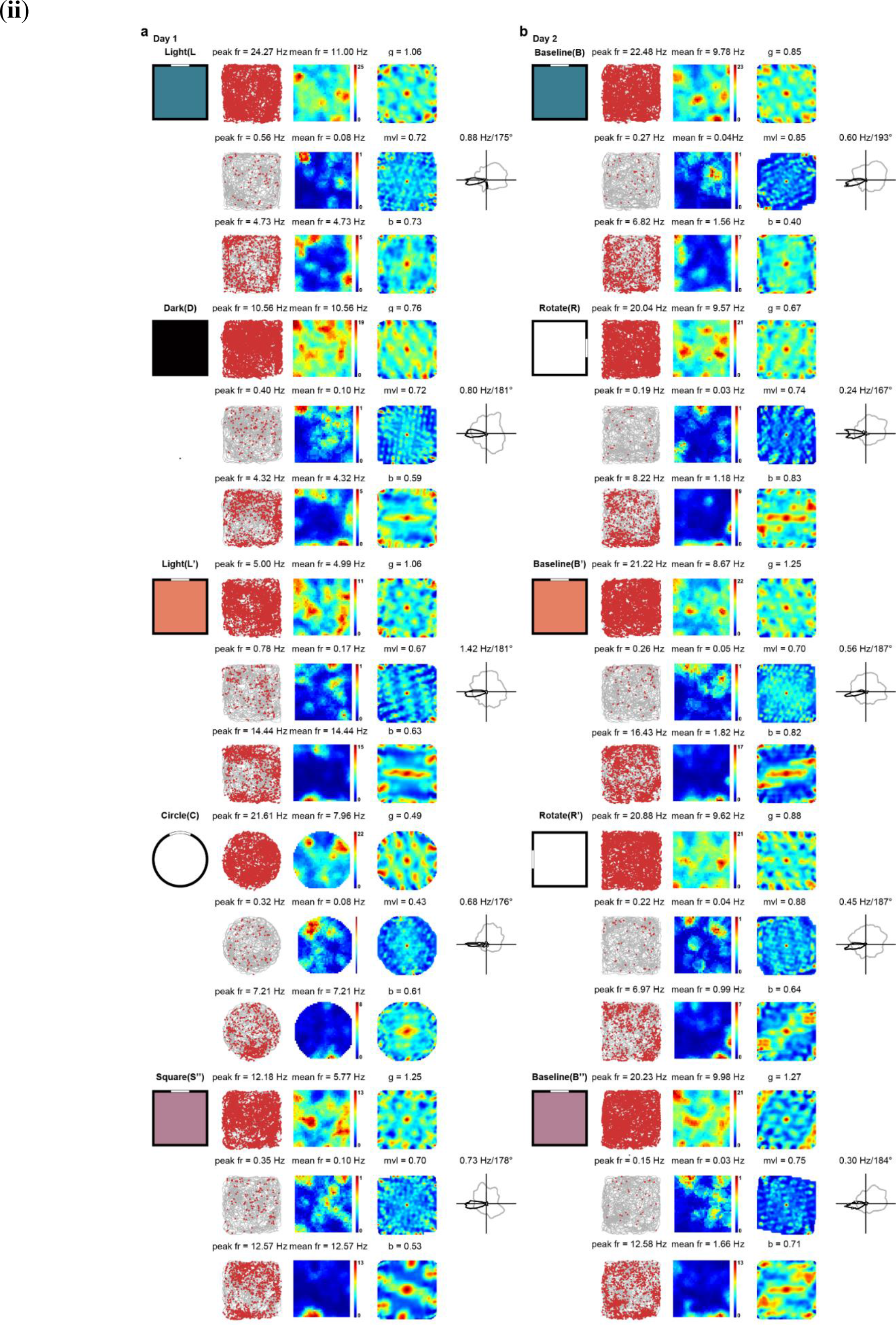
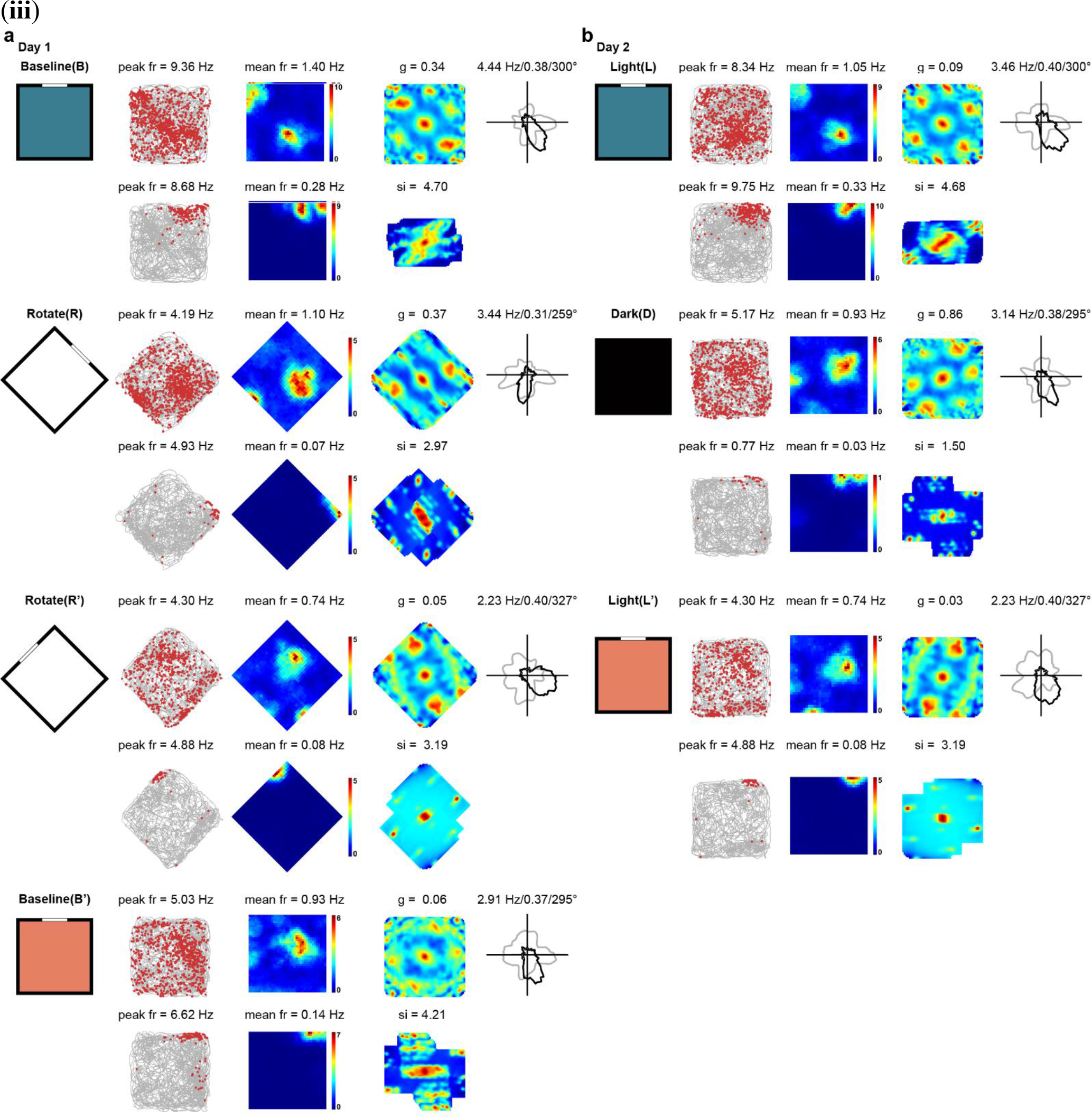
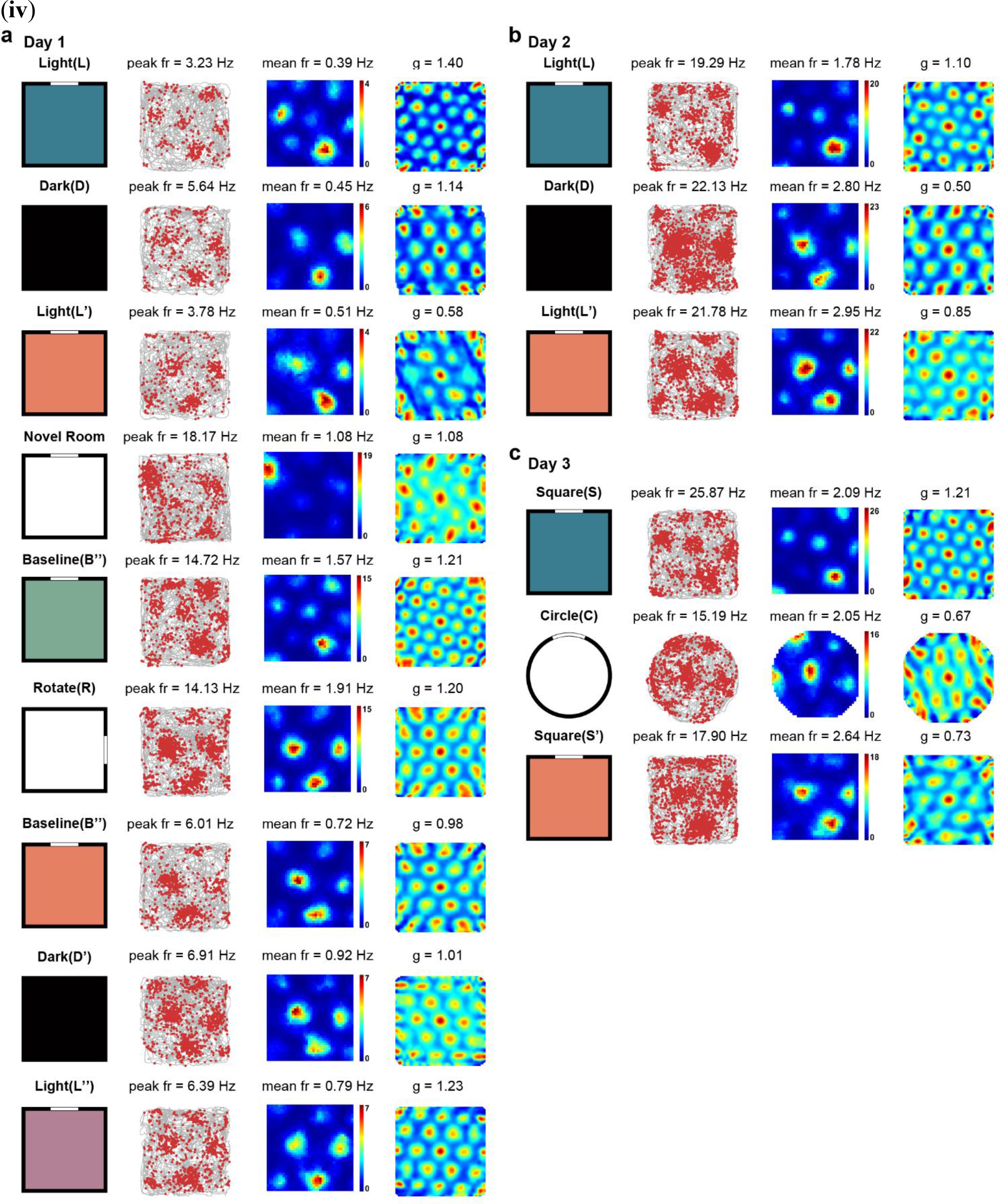
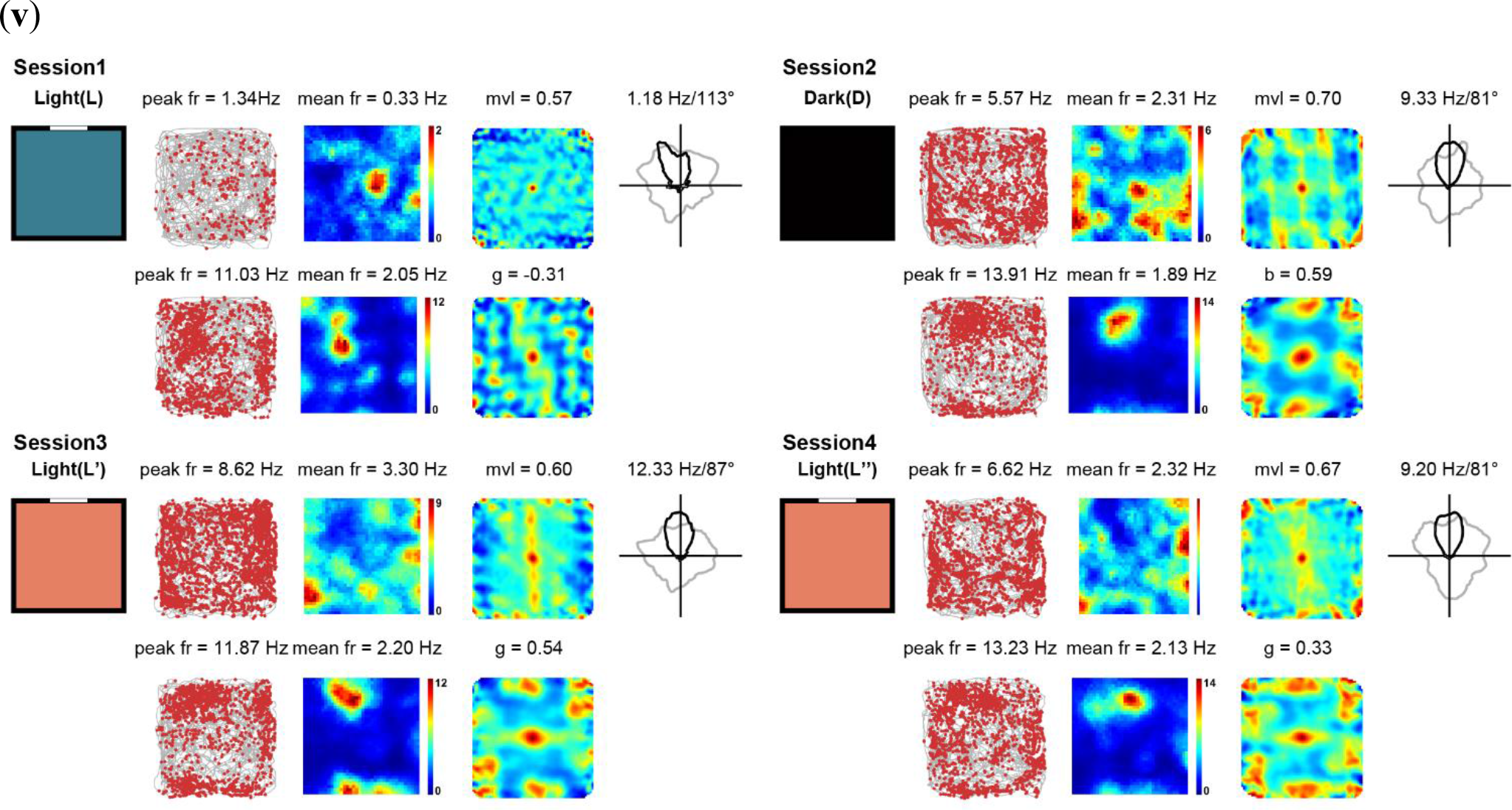
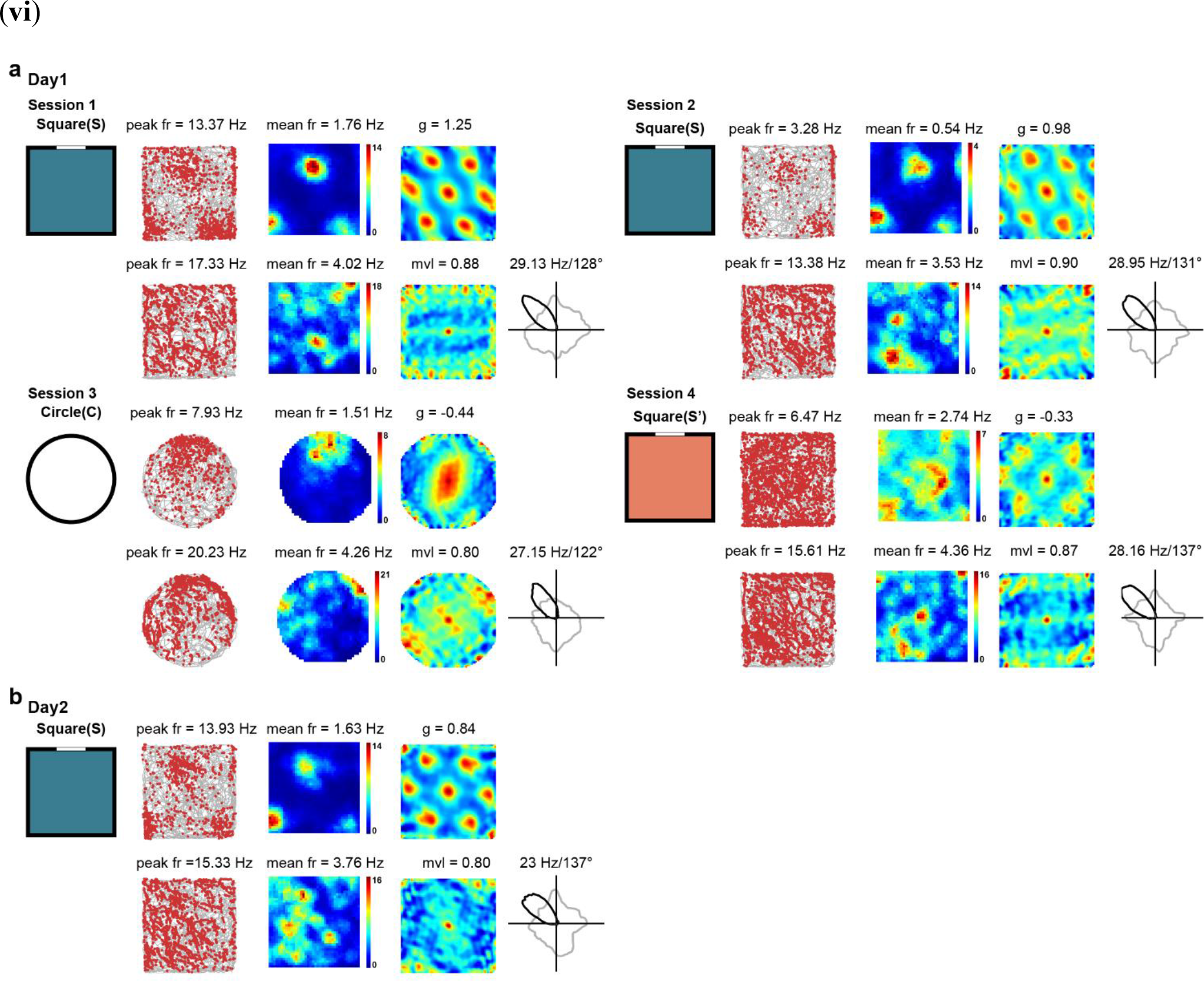
Full sample of bimodal grid cells in V2. (i-vi) Switch of grid firing patterns in responses to changes in a familiar environment and restoration of the grid pattern after re-entry into the same environment from a novel room or from animals’ home cage. (i) **(a)** Simultaneously recording of one head-direction cell, one bimodal grid cell and one border cell. **(b)** Simultaneously recording of two bimodal grid cells and one border cell. (ii) **(a)** Simultaneously recording of one bimodal grid cell, one head-direction cell and one border cell in V2 on day 1. **(b)** Simultaneously recording of same three spatial cells on day 2. (iii) **(a)** Simultaneously recording of one conjunctive bimodal grid cell and one place cell in V2 on day 1. **(b)** Simultaneously recording of same two spatial cells on day 2. (**iv**) **(a)** Bimodal switch of a bimodal grid cell in responses to environmental changes and restoration of the original pattern after re-entry into the same familiar environment. **(b)** Bimodal switch of grid firing patterns after total darkness. **(c)** Recording of the same bimodal grid cell across different shapes of running boxes. (**v**) Simultaneously recording of one head-direction cell and one bimodal grid cell in V2 across light-light- dark-light sessions. (**vi**) **(a)** Simultaneously recording of one bimodal grid cell and one head-direction cell. **(b)** Simultaneously recording of same two spatial cells in **(a)** on day 2.

**Fig. S10.**
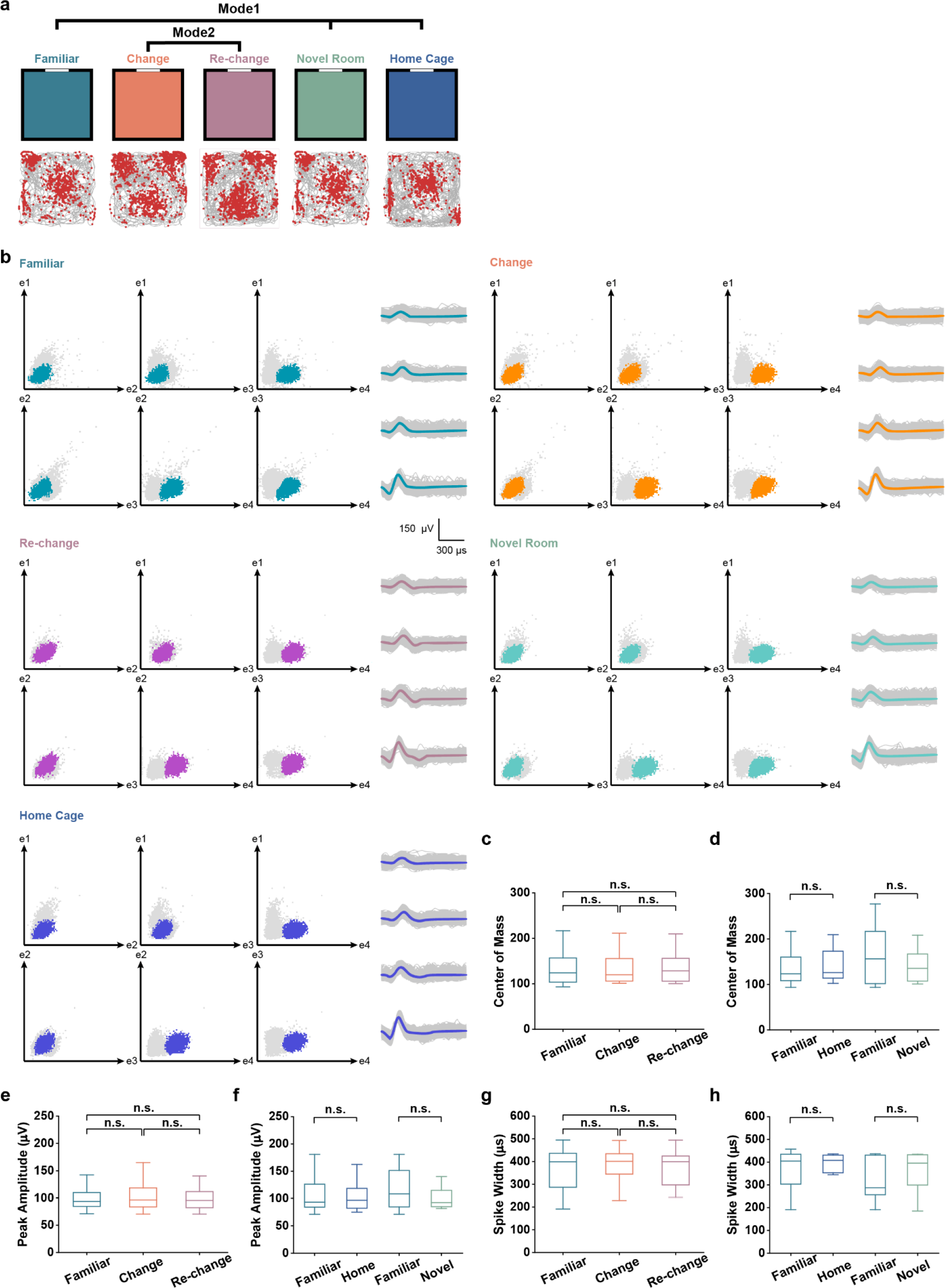
Persistence of cluster diagram and spike waveform during bimodal switch of V2 grid firing patterns. (a) Bimodal firing patterns of the same grid cell in **Fig. 2** under familiar environment, after environmental change, re-change, re-entry from a novel room and re-entry from the home cage. (b) Cluster diagram (left panel) showing the peak to trough amplitudes of recorded spikes on each pair of electrodes (e1-e4) on a tetrode. Each dot represents a single spike with the color dots indicating the spike cluster for the grid cell showing in (**a**). Waveforms on each of the four electrodes for the gird cells are shown on the right panel. Spike clusters and waveforms are preserved during bimodal switch of grid firing patterns. (**c**-**d**) Centers of mass for the spike clusters of V2 bimodal grid cells are stable during shift (two-sided Wilcoxon signed-rank test, Familiar-Change: *Z* = 0.23, *P* = 0.82; Change-Rechange: *Z* = 1.36, *P* = 0.17; Familiar-Rechange, *Z* = 0.45, *P* = 0.65, *n* = 15) and restoration (Familiar-Home cage: *Z* = 0.36, *P* = 0.72, *n* = 12; Familiar-Novel room: *Z* = 0.85, *P* = 0.40, *n* = 7) of grid structures. (**e**-**f**) Peak amplitudes of V2 bimodal grid cells remain unaltered during shift (Familiar-Change: *Z* = 0.31, *P* = 0.76; Change-Rechange: *Z* = 1.19, *P* = 0.23; Familiar-Rechange, *Z* = 0.43, *P* = 0.67) and restoration (Familiar-Home cage: *Z* = 0.94, *P* = 0.35; Familiar-Novel room: *Z* = 1.01, *P* = 0.31) of grid structures. (**g**-**h**) Spike widths of V2 bimodal grid cells maintain during shift (Familiar-Change: *Z* = 0.82, *P* = 0.41; Change-Rechange: *Z* = 1.36, *P* = 0.17; Familiar-Rechange, *Z* = 0.51, *P* = 0.61) and restoration (Familiar- Home cage: *Z* = 0.82, *P* = 0.41; Familiar-Novel room: *Z* = 0.85, *P* = 0.40) of grid firing patterns.

